# SMCHD1’s DNA binding activity enables its stable retention on chromatin

**DOI:** 10.64898/2026.09.14.751594

**Authors:** Ruifeng Hu, Julissa Sanchez-Velasquez, Jieqiong Lou, Kelsey Breslin, Tamara Cameron, Ashleigh N. Solano, Iromi Wickramasinghe, Quentin Gouil, Andrew Kueh, Tracy Willson, Andrew Keniry, Niall D. Geoghegan, Elizabeth Hinde, Marnie Blewitt

## Abstract

Chromatin proteins play critical roles in gene regulation, yet frequently we do not fully understand how weak DNA binding affinity of such proteins contributes to their locus-specific actions. Here, we studied SMCHD1, a non-canonical SMC-family protein involved in three-dimensional genome organization and gene repression of the inactive X chromosome and its autosomal targets. We replaced endogenous SMCHD1 with GFP-tagged wild-type or hinge-domain DNA-binding mutant SMCHD1 to define the cellular role of DNA binding. The mutant showed reduced enrichment at the inactive X chromosome in female cells, while retaining stable binding at most autosomal binding sites. Impaired DNA binding weakens SMCHD1-mediated gene repression and chromatin-state regulation, producing hypomorphic effect. Multiple live-cell imaging methods reveal that DNA binding constrains SMCHD1 mobility and supports maintenance, rather than initial recruitment, of chromatin-bound SMCHD1 both during interphase and mitosis. Thus, SMCHD1’s weak and sequence-independent DNA binding is a key determinant of its chromatin residence, localization and function. Our findings provide a framework for understanding SMCHD1 and other chromatin proteins with sequence-independent DNA binding activity.

## Main

Transcription factors decode cis-regulatory information through sequence-specific DNA binding, forming a core component of the gene-regulatory code^1, 2^. This binding provides a molecular basis for transcription factor-mediated gene regulation by controlling polymerase recruitment, chromatin state and higher-order genome organization^3–5^. By contrast, many epigenetic regulators have essential roles in gene regulation despite lacking intrinsic sequence-specific DNA-binding activity. Their function therefore depends on chromatin association, chromatin residence and interactions with other chromatin-bound factors^6^. Recent work also indicates that weak DNA-binding events can stabilize chromatin interactions mediated by intrinsically disordered regions, converting dynamic chromatin sampling into reproducible occupancy at specific genomic regions^7^. Thus, DNA interaction and chromatin interaction provide complementary mechanisms that enable regulatory proteins to search, recognize and remain associated with their genomic targets^8, 9^. However, how DNA binding shapes chromatin residence and supports the function of epigenetic regulators lacking sequence-specific DNA binding remains poorly understood.

Beyond static DNA or chromatin association, the dynamic behaviour of regulatory proteins in living cells is central to their function. For transcription factors, chromatin residence time can vary at different targets independently of protein concentration but be affected by mutations or post-translational modifications. Prolonged chromatin residence is often associated with stronger transcriptional activation or repression^10–12^. Altered residence time can also affect linked molecular processes; for example, reduced chromatin residence of mutant CTCF disrupts three-dimensional genome organization and DNA methylation patterns^13^. Quantitative analysis of residence time and molecular mobility therefore provides an important framework for understanding epigenetic regulator function in living cells.

Structural Maintenance of Chromosomes Hinge Domain Containing 1 (SMCHD1) is a non-canonical SMC-family epigenetic regulator involved in gene repression, chromatin state and three-dimensional genome organization^14–20^. SMCHD1 contributes to repression of multiple clustered gene families or regions, including protocadherin and *Hox* gene clusters, imprinted gene clusters and the inactive X chromosome (Xi) in females^16, 21, 22^. Acute somatic deletion of SMCHD1 only partially derepresses some of its targets, notably not reactivating the Xi, whereas loss during early development and differentiation causes more extensive gene activation, suggesting that SMCHD1’s primary role is during the establishment of silencing and is context-dependent^17, 23–25^.

SMCHD1 shapes repressive chromatin landscapes by supporting acquisition of CpG island DNA methylation and H3K9me3-marked heterochromatin at both autosomal targets and the Xi^15, 20, 25^. Its interplay with repressive chromatin marks is particularly evident at the Xi, where SMCHD1 is recruited downstream of the polycomb repressive complex 1 (PRC1) mark H2AK119ub and accumulates across the Xi territory^24, 26^. Loss of SMCHD1 generally increases PRC2 mark H3K27me3 spreading across the Xi, while reducing H3K27me3 at selected gene-proximal regions prone to derepression, indicating that SMCHD1 fine-tunes heterochromatin distribution rather than simply promoting or opposing a single repressive state^18, 27^. SMCHD1 also contributes to the distinctive three-dimensional architecture of the Xi by antagonizing topologically associating domain formation and promoting long-range chromatin interactions, partly through regulation of CTCF occupancy^17–20^.

SMCHD1 contains an N-terminal ATPase domain with neighbouring ubiquitin-like (UBL) and bromo-adjacent (BAH) domains, a poorly characterized central region that contains at least one immunoglobulin-like (IGL) domain followed by a C-terminal hinge domain surrounded by coiled-coil domains^28^. The ATPase domain is a GHKL ATPase type and hydrolyses ATP, a process required for proper SMCHD1 localization and function at pericentric heterochromatin and the Xi^29–31^. The hinge domain is related to that of canonical SMC proteins and mediates homodimerization and DNA binding^32, 33^. Recent work from our group showed that the BAH and IGL domains can also interact with DNA, and that DNA binding stimulates ATPase activity in vitro^31^, suggesting SMCHD1 engages with DNA in a multivalent fashion. However, SMCHD1 does not appear to show strong sequence-specific DNA-binding preference in vivo^16^. SMCHD1 is essential for normal development^14^, and pathogenic variants in human SMCHD1 are associated with facioscapulohumeral muscular dystrophy (FSHD) and Bosma arhinia microphthalmia syndrome (BAMS)^34–36^. Several disease-associated variants lie within the characterised domains and alter SMCHD1 ATPase activity or DNA binding in vitro. While recent work has studied how the ATPase activity relates to function in cells^30, 31^, nothing is understood about how the DNA binding activity of the hinge domain shapes SMCHD1’s molecular function.

Although many pathogenic variants reduce SMCHD1 abundance, some patient-derived and engineered variants alter SMCHD1 function without detectably changing protein levels, suggesting that additional biochemical properties contribute to SMCHD1 activity independently of protein abundance^30, 34, 35, 37^. Here, we focus on the FSHD-associated hinge-domain variant R1866G (R1867G in mouse), which impairs SMCHD1 DNA-binding activity in vitro^16, 33^. Combining genomics with live-cell imaging, we show that DNA binding does not act as a primary recruitment mechanism, but instead constrains SMCHD1 mobility, prolongs chromatin residence and enables full SMCHD1 function.

## Results

### DNA binding enables SMCHD1 enrichment at the inactive X chromosome

To define the contribution of DNA binding of the hinge domain to SMCHD1 function, we generated a switchable mouse model carrying the well-characterized FSHD-associated SMCHD1 R1867G variant (R1866G in human SMCHD1)^38^. We inserted a LoxP-STOP-LoxP (LSL) cassette followed by cDNA encoding full-length, C-terminally GFP-tagged wild-type SMCHD1 (WT) or SMCHD1 R1867G into the *Rosa26* locus. Crossing these mice with our existing *Smchd1* conditional-knockout strain^17^, and the *Rosa26-CreERT2* line^39^ generated a scenario in which 4-hydroxytamoxifen (4-OHT) treatment simultaneously deleted endogenous *Smchd1* and induced either WT or R1867G SMCHD1-GFP (Fig. 1a). We derived mouse embryonic fibroblasts (MEFs) and neural stem cells (NSCs) from embryonic day 14.5 embryos and treated the primary cells with 4-OHT for 1-5 days. Five days of treatment were sufficient to complete the switch in SMCHD1 protein expression (Extended Data Fig. 1a, b). Importantly, the WT and R1867G proteins were expressed at similar levels (Extended Data Fig. 1c).

**Fig. 1.**
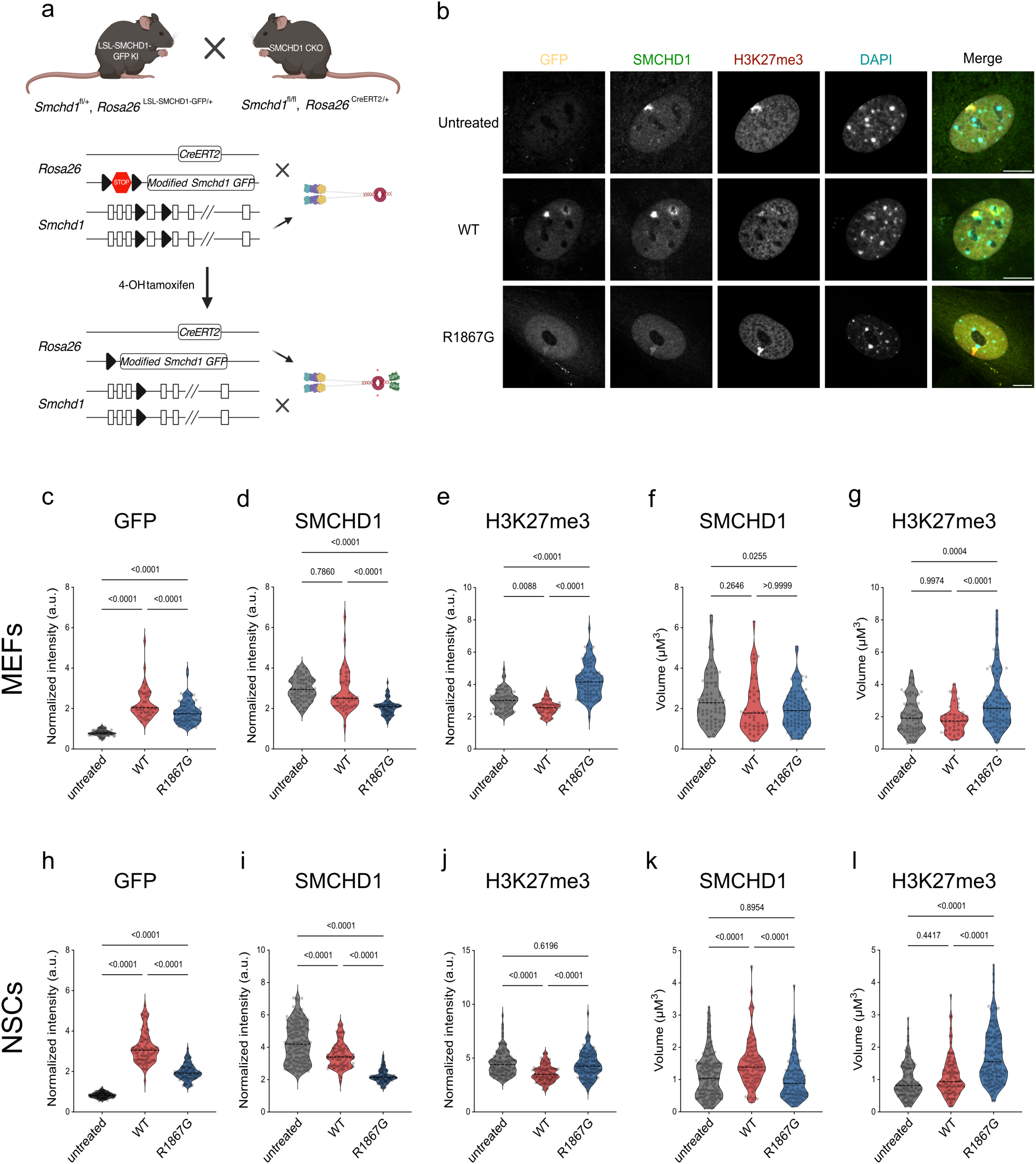
Generation of a switchable SMCHD1 model reveals that DNA binding is required for SMCHD1 enrichment at the Xi. a, Schematic of the mouse model used to generate switchable SMCHD1 cell lines expressing GFP-tagged WT or R1867G SMCHD1. b, Representative immunofluorescence images of switchable WT and R1867G MEFs after 4-OHT treatment. Scale bar, 10 µm. c-e, Normalized mean GFP, SMCHD1 and H3K27me3 fluorescence intensity, respectively, within the Xi territory of MEFs. f, g, Three-dimensional volume of the SMCHD1- and H3K27me3-enriched Xi domains, respectively, in MEFs. n = 66 untreated, 42 WT and 82 R1867G cells. h-j, Normalized mean GFP, SMCHD1 and H3K27me3 fluorescence intensity, respectively, within the Xi territory of NSCs. k, l, Three-dimensional volume of the SMCHD1- and H3K27me3-enriched Xi domains, respectively, in NSCs. n = 150 untreated, 79 WT and 100 R1867G cells. Data were pooled from three independent experiments. P values were calculated by one-way ANOVA with Bonferroni correction for multiple comparisons.

Next, we asked how the R1867G variant altered the Xi by immunofluorescence staining for GFP, SMCHD1 and the Xi-associated mark H3K27me3. This confirmed that induced WT SMCHD1-GFP was recruited to the Xi (Extended Data Fig. 1d). Compared with WT MEFs, R1867G MEFs less frequently displayed a detectable SMCHD1 Xi focus, but more frequently displayed an H3K27me3-enriched Xi domain (Extended Data Fig. 1e, g-j), suggesting that impaired DNA binding compromises SMCHD1 accumulation at the Xi (Fig. 1b). Because H3K27me3 spreading on the Xi is altered by loss or mutation of SMCHD1^17, 25, 31, 37^, we independently segmented the Xi territory using either SMCHD1 or H3K27me3 signal and quantified the corresponding fluorescence intensity and volume. To enable direct comparison of the two domains, this analysis was restricted to MEFs with detectable SMCHD1 and H3K27me3 Xi signals. R1867G MEFs showed reduced GFP and SMCHD1 intensity at the Xi, together with increased H3K27me3 intensity and an expanded H3K27me3-enriched volume (Fig. 1c-e, g). The volume of the detectable SMCHD1 Xi domain did not differ between WT and R1867G MEFs (Fig. 1f). However, requiring a detectable SMCHD1 Xi focus probably enriched the R1867G group for MEFs retaining relatively strong Xi association and may therefore underestimate the magnitude of the R1867G defect.

Quantitative immunofluorescence analysis in NSCs yielded similar results. Most R1867G NSCs retained a detectable SMCHD1 Xi focus, although the fraction of cells displaying an H3K27me3-enriched Xi domain was increased relative to WT cells (Extended Data Fig. 1f, k-n). Within individual Xi territories, R1867G NSCs showed reduced GFP and SMCHD1 intensity and a smaller SMCHD1-enriched volume (Fig. 1h, i, k). Conversely, H3K27me3 intensity and Xi-domain volume were increased (Fig. 1j, l). Together, these findings establish that hinge-domain DNA binding is required for robust SMCHD1 accumulation at the Xi and show that reduced SMCHD1 occupancy is accompanied by expansion of the H3K27me3-enriched Xi compartment, consistent with prior data^17, 25, 37^.

### Impaired DNA binding has limited effects on SMCHD1 occupancy outside the Xi

To determine how DNA binding affects genome-wide SMCHD1 occupancy, we performed chromatin immunoprecipitation followed by sequencing (ChIP-seq) for the GFP tag in the wild-type and R1867G variant female NSCs (n=3 per genotype) and compared to a control that had no GFP tag (GFPneg). Before genomic profiling, we examined the protein level of induced SMCHD1-GFP in individual cell lines and confirmed similar protein level after 4-OHT treatment (Extended Data Fig. 2a). Using MACS peak calling^40^, we identified 472 significant peaks of SMCHD1 binding in the WT-expressing samples (29 ChrX peaks) and 502 peaks in the R1867G-expressing samples (24 ChrX peaks) (Supplementary file 1). Because this peak calling analysis only reveals limited narrow strong binding regions, we also investigated broader enrichment patterns. When normalising ChIP signals to the corresponding input signals, we found that only the X chromosome, and not the autosomes, showed chromosome-wide SMCHD1 signal enrichment (Figure. 2a, b, Extended Data Fig. 2b), suggesting this signal is Xi derived. ChIP-seq revealed a broad reduction in SMCHD1 signal across almost the entire X chromosome in R1867G cells (Fig. 2a, b). The switchable cells lacked the required genetics to distinguish the Xi from the active X chromosome (Xa). However, previous studies showed that SMCHD1 is enriched across the Xi but not Xa in female cells^17–19^. We therefore interpret the chromosome-wide reduction in X-linked SMCHD1 signal in female samples as a loss of occupancy across the Xi, consistent with the immunofluorescence data.

**Fig. 2.**
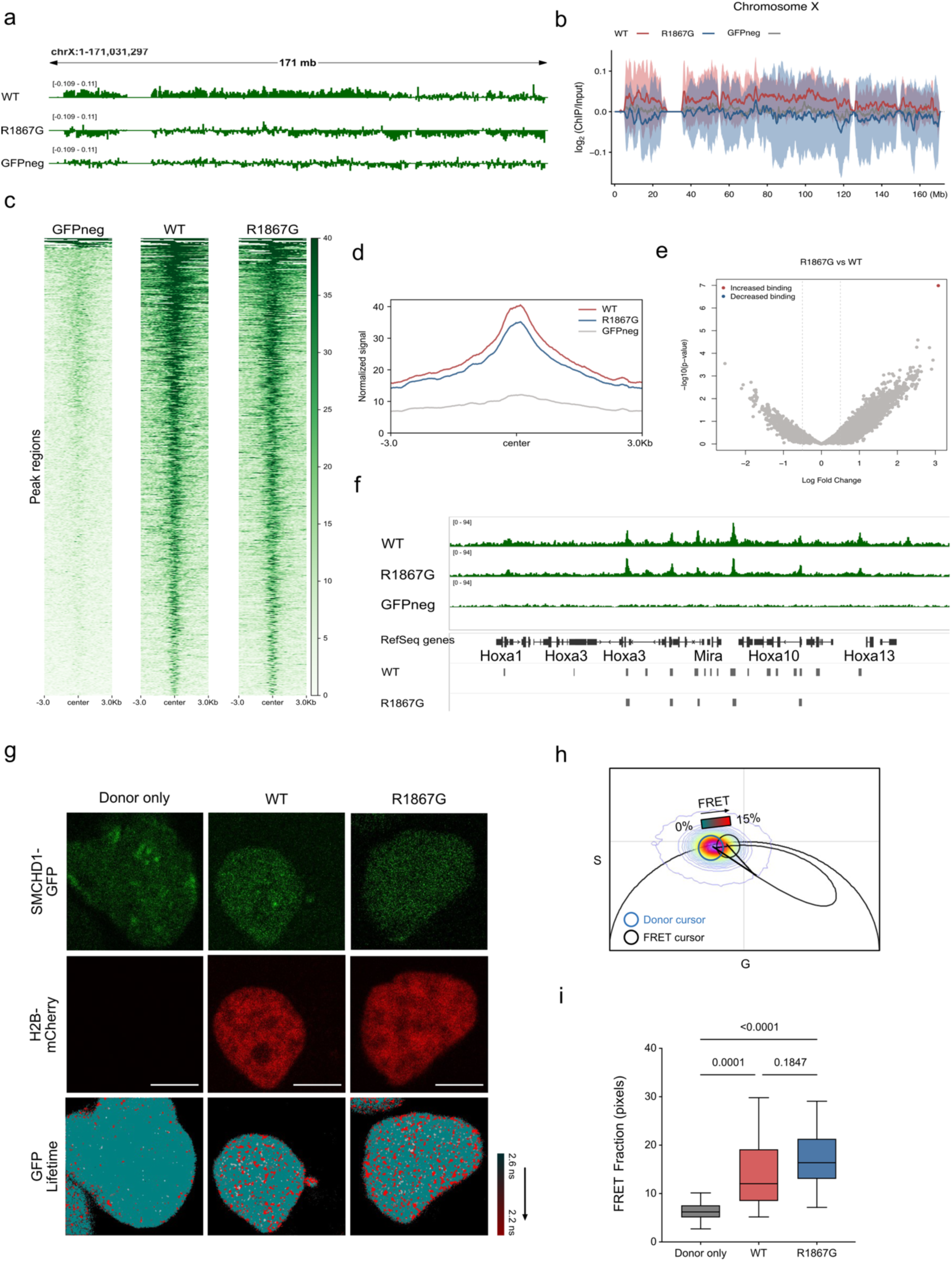
Impaired DNA binding preferentially reduces SMCHD1 occupancy across the X chromosome. a, Chromosome-wide profiles of WT and R1867G SMCHD1 occupancy across the X chromosome. ChIP-seq signal was normalized to the corresponding input and is shown as log2(ChIP/input). Tracks show the mean of biological replicates (WT, n = 3; R1867G, n = 3; GFP-negative control, n = 1). Bin size, 50 kb. b, Aggregate SMCHD1 occupancy across the X chromosome. Solid lines indicate the mean and shaded regions indicate 95% confidence intervals. Bin size, 1 Mb. c, d, Spike-in-normalized SMCHD1 signal across reproducible MACS-called peaks and flanking regions. e, Differential SMCHD1 occupancy between WT and R1867G samples, analysed using csaw package (FDR < 0.05). f, SMCHD1 occupancy at the *HoxA* locus, with corresponding MACS-called peaks. g, Representative images show interaction between SMCHD1 and histone H2B in live male NSCs. Top panel: NSCs expressing SMCHD1-GFP; Middle panel: NSCs transfected with or without H2B-mCherry; Bottom panel: pseudocolored GFP lifetime maps of cells. Scale bar, 5 µm. h, Phasor distribution of GFP fluorescence lifetime from donor only and FRET sample shown in g with the theoretical FRET trajectory superimposed to determine the range of FRET efficiencies. i, Fraction of pixels that detected FRET signal of each group. Combined samples of two independent experiments. n = 16 Donor only, 25 WT and 21 R1867G cells. P values were calculated by one-way ANOVA with Bonferroni correction for multiple comparisons.

SMCHD1 also occupies discrete autosomal regions and regulates autosomal gene clusters^16, 17, 37^. After spike-in normalisation, SMCHD1 signal at autosomal peaks and their flanking regions was largely preserved in R1867G cells (Fig. 2c, d). Differential-binding analysis identified only one region with significantly increased SMCHD1 occupancy in R1867G cells (Fig. 2e, Extended Data Fig. 2c), while prominent autosomal targets, including the *HoxA* locus, retained clear SMCHD1 binding (Fig. 2f). Thus, the R1867G substitution preferentially compromises the broad Xi-associated SMCHD1 domain while leaving high-occupancy autosomal binding sites largely intact. Our ChIP-seq protocol was optimised to resolve the broad SMCHD1 domain across the Xi and was comparatively stringent (Methods). We used ionic-detergent-free buffer to avoid disruption of SMCHD1 and prolonged sonication time to 30 mins to reveal SMCHD1 enrichment at compacted chromatin regions such as the inactive X chromosome. It therefore preferentially detected regions of high or stable SMCHD1 occupancy relative to less stringent ChIP-seq approaches (Extended Data Fig. 2d)^37^. We cannot exclude the possibility that R1867G alters transient or low-occupancy SMCHD1 interactions that were not captured by this assay.

To further examine how impaired DNA binding affects SMCHD1 association with chromatin, we employed fluorescence lifetime imaging microscopy coupled with Förster resonance energy transfer (FLIM-FRET) to measure SMCHD1 proximity to histone H2B, within approximately 10 nm in living cells^41, 42^. This involved transiently expressing mCherry-tagged histone H2B in female NSCs as a FRET acceptor and measuring changes in the fluorescence lifetime of SMCHD1-GFP as the donor. FRET, detected as a reduced SMCHD1-GFP donor lifetime, was readily observed in cells expressing H2B-mCherry, indicating that SMCHD1 interacts with chromatin in living cells (Extended Data Fig. 2e). Notably, WT and R1867G SMCHD1 showed comparable whole-nucleus FRET signals (Extended Data Fig. 2f), suggesting that global SMCHD1 proximity to chromatin is not disrupted by impaired hinge-domain DNA binding. In female NSCs, however, SMCHD1-GFP was markedly enriched at the Xi territory compared with the rest of the nucleus, resulting in a high local donor-to-acceptor ratio that compromised reliable FLIM-FRET detection^43^ (Extended Data Fig. 2g). Thus, to avoid this bias, we also performed FLIM-FRET in male NSCs, in which SMCHD1-GFP showed a more uniform nuclear distribution (Fig. 2g). In male NSCs, FRET was also readily detected in cells expressing H2B-mCherry (Fig. 2g, h). Importantly, WT and R1867G SMCHD1 again showed comparable whole-nucleus FRET signals (Fig. 2i), suggesting that impaired hinge-domain DNA binding does not markedly disrupt SMCHD1 proximity to chromatin outside the Xi context. Together with the ChIP-seq results, these findings indicate that hinge-domain DNA binding is particularly important for chromosome-wide SMCHD1 accumulation at the Xi, whereas stable chromatin association at autosomal regions is largely retained.

### Impaired DNA binding produces a hypomorphic SMCHD1 state

To assess the functional consequences of impaired hinge-domain DNA binding, we performed RNA sequencing (RNA-seq). In addition to our switchable *Smchd1* cells, we included conditional *Smchd1*-knockout NSCs (*Smchd1*^fl/fl^; *Rosa26*^CreERT2/+,^ hereafter KO) and matched control NSCs (*Smchd1*^fl/fl^; *Rosa26*^+/+^, hereafter CON), enabling comparison of the R1867G phenotype with complete SMCHD1 loss. Multidimensional scaling separated samples according to genotype (Extended Data Fig. 3a). Comparison of KO and CON NSCs identified 790 differentially expressed genes, including established SMCHD1-sensitive loci such as the protocadherin (*Pcdh*) clusters^16, 21, 22^ (Fig. 3a, Supplementary file 2). We did not detect activation of X-linked genes in KO NSCs (Fig. 3a), consistent with previous evidence that post-X-inactivation loss of SMCHD1 is insufficient to reactivate genes silenced on the Xi^17, 19, 24, 27^. By contrast, no individual gene met the genome-wide differential-expression threshold in R1867G versus WT NSCs (Fig. 3b). We therefore separated the genes upregulated or downregulated in KO cells into two SMCHD1-sensitive gene sets and tested their behaviour in R1867G cells. Both gene sets significantly shifted concordantly in R1867G cells: genes upregulated in KO cells tended to increase, whereas genes downregulated in KO cells tended to decrease (ROAST, FDR_up_ = 0.0035 and FDR_down_ = 0.011; Fig. 3c). The *Pcdh* genes upregulated in KO cells similarly showed intermediate increases in R1867G cells relative to WT cells (Extended Data Fig. 3b). *Smchd1* transcript abundance was comparable between WT and R1867G cells (Extended Data Fig. 3c), consistent with the comparable protein abundance observed by immunoblotting (Extended Data Fig. 2a). Thus, R1867G produces a partial, SMCHD1-loss-like transcriptional phenotype that reflects impaired protein function rather than altered SMCHD1 abundance.

**Fig. 3.**
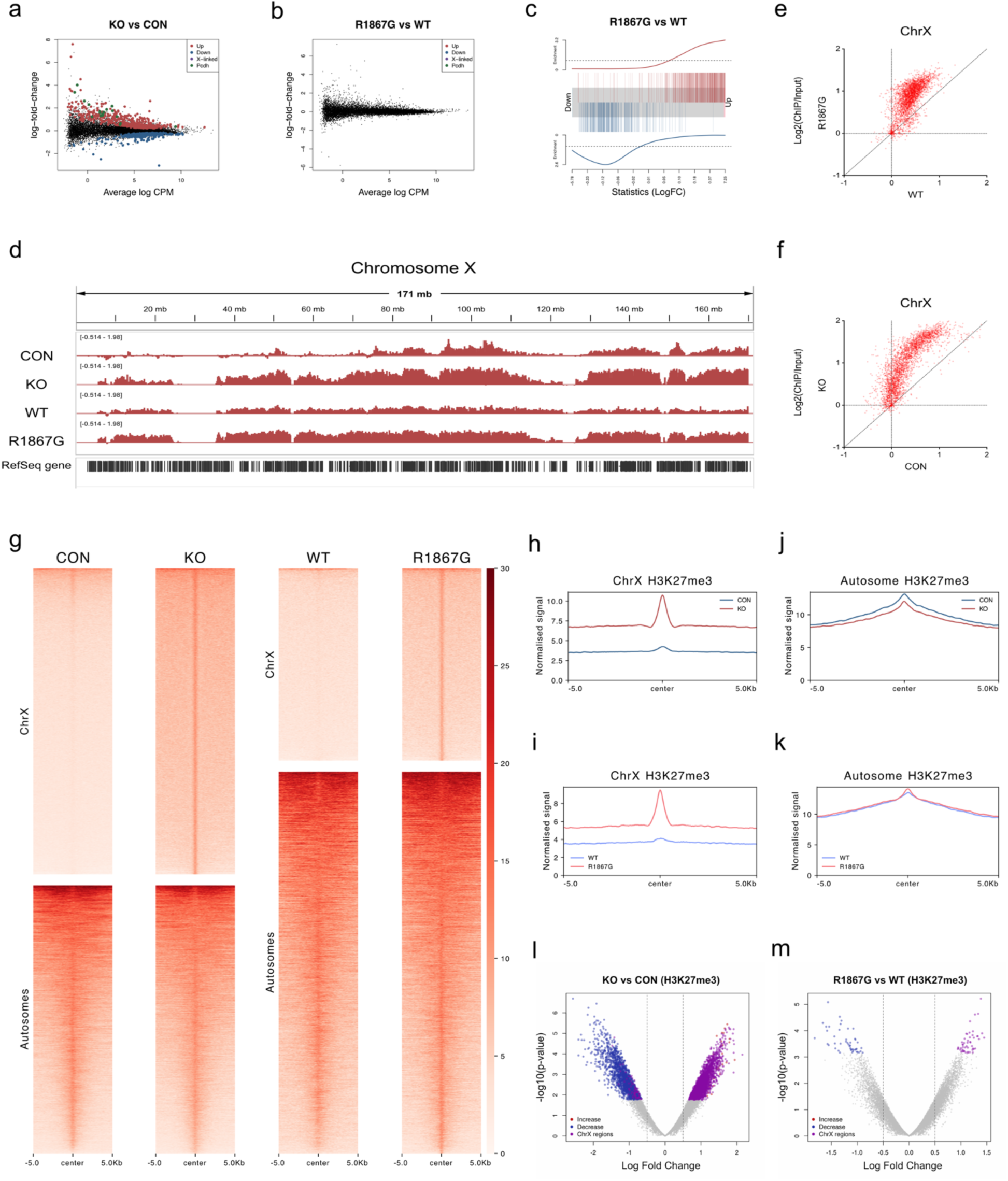
Impaired DNA binding partially phenocopies SMCHD1 loss at the transcriptome and H3K27me3 landscape. a, b, MA plots showing differential gene expression in KO versus CON NSCs (a) and switched R1867G versus WT female NSCs (b), analysed using edgeR (FDR < 0.05). c, Barcode plots showing expression changes in R1867G cells among genes identified as differentially expressed in SMCHD1-KO cells (ROAST; FDRup = 0.0035 and FDRdown = 0.011). d, Mean H3K27me3 ChIP-seq signal across the X chromosome in CON, KO, WT and R1867G NSCs, shown as log2(ChIP/input). Biological replicates: CON and KO, n = 2 each; WT and R1867G, n = 3 each. e,f, Paired comparison of H3K27me3 signal in 50-kb bins across the X chromosome. g, Spike-in-normalized mean H3K27me3 signal, expressed as reads per kilobase per million mapped reads (RPKM), across reproducible peak regions on autosomes and the X chromosome. h-k, Line plots of the H3K27me3 profiles summarized in g. l, Differential H3K27me3 enrichment in KO versus CON cells, analysed using csaw (FDR < 0.05). Differentially enriched regions are coloured; purple denotes regions on the X chromosome. m, Differential H3K27me3 enrichment in R1867G versus WT switched cells, analysed using csaw (FDR < 0.1).

We next profiled H3K27me3, which showed altered association to the Xi following SMCHD1 switching immunofluorescence (Fig. 1). By ChIP-seq, SMCHD1 loss increased H3K27me3 signal across the Xi and expanded the H3K27me3-enriched domain (Fig. 3d, e). As H3K27me3 is broadly enriched across the Xi but not Xa in female cells^44, 45^, the X-chromosome profiles likely reflect Xi-associated signal. The H3K27me3 profile in switched WT cells did not fully recapitulate that of unswitched control cells, displaying a more even distribution and broader enriched regions (Fig. 3d). This may reflect the transition from endogenous SMCHD1 depletion to SMCHD1-GFP induction during the switching procedure. We therefore compared R1867G and WT cells subjected to the same switching regimen. Compared to its matched switched control, R1867G cells displayed increased H3K27me3 signal across most of the X chromosome, recapitulating the direction of change observed after SMCHD1 loss (Fig. 3d, f).

After spike-in normalization, peak-based analysis further showed increased H3K27me3 signal at Xi-associated peaks in both KO and R1867G cells (Fig. 3g-i). At autosomal peaks, SMCHD1 loss slightly reduced H3K27me3 signal, whereas R1867G had only very minor effects (Fig. 3j, k). Differential-enrichment analysis identified thousands of H3K27me3 regions altered in KO cells: most regions with decreased signal were autosomal, whereas most regions with increased signal mapped to the X chromosome (Fig. 3l, Supplementary file 3,4). These data extend previous observations of context-dependent H3K27me3 changes at the Xi after SMCHD1 loss^18, 27^, suggest that SMCHD1 influences H3K27me3 through distinct mechanisms at the Xi and autosomal loci (Extended Data Fig. 3d). R1867G cells contained fewer differentially enriched regions, but changes followed the same overall direction when analysed at a relaxed statistical threshold (Fig. 3m).

We next asked whether these regional changes reflected altered global H3K27me3 abundance. Immunoblotting showed no detectable change in total H3K27me3 in KO or R1867G NSCs (Extended Data Fig. 3e). Re-analysis of the quantitative immunofluorescence data similarly showed comparable total nuclear H3K27me3 intensity in WT and R1867G NSCs, despite increased H3K27me3 signal within the Xi (Extended Data Fig. 3f, g). H3K27me3 intensity in the remainder of the nucleus was unchanged (Extended Data Fig. 3h), probably because the Xi-associated pool represented only a small fraction of total nuclear H3K27me3 (Extended Data Fig. 3i). Thus, impaired DNA binding redistributes H3K27me3 at specific chromosomal domains without measurably altering its global abundance. Together, the transcriptional and chromatin profiles define R1867G as a hypomorphic SMCHD1 variant and demonstrate that hinge-domain DNA binding is required for full SMCHD1 function.

### DNA binding prolongs SMCHD1 chromatin residence without controlling initial recruitment

We next investigated how impaired hinge-domain DNA binding compromises SMCHD1 residence time on the chromatin. We hypothesized that weaker DNA interactions reduce SMCHD1 residence time on chromatin. Chromatin residence time, independently of protein abundance, is an important determinant of transcription-factor and chromatin-regulator activity, as exemplified by CTCF^13^. To measure SMCHD1 dynamics in living cells, we performed fluorescence recovery after photobleaching (FRAP) at the Xi in female MEFs (Fig. 4a). Because SMCHD1 is highly concentrated at the Xi and R1867G reduces this enrichment, we also measured nucleoplasmic dynamics in male MEFs, in which WT and R1867G cells contained comparable total nuclear SMCHD1-GFP levels (Fig. 4a, Extended Data Fig. 4b). This design minimized potential effects of unequal partitioning of the protein between the Xi and the nucleoplasm. Consistent with previous measurements in mouse embryonic stem cells^30^, SMCHD1 recovered slowly at the Xi but rapidly in the nucleoplasm (Fig. 4a). Nucleoplasmic fluorescence approached a plateau within approximately 1 min (Extended Data Fig. 4a and Supplementary Video 1), suggesting that diffusion makes a major contribution to recovery in this compartment.

**Fig. 4.**
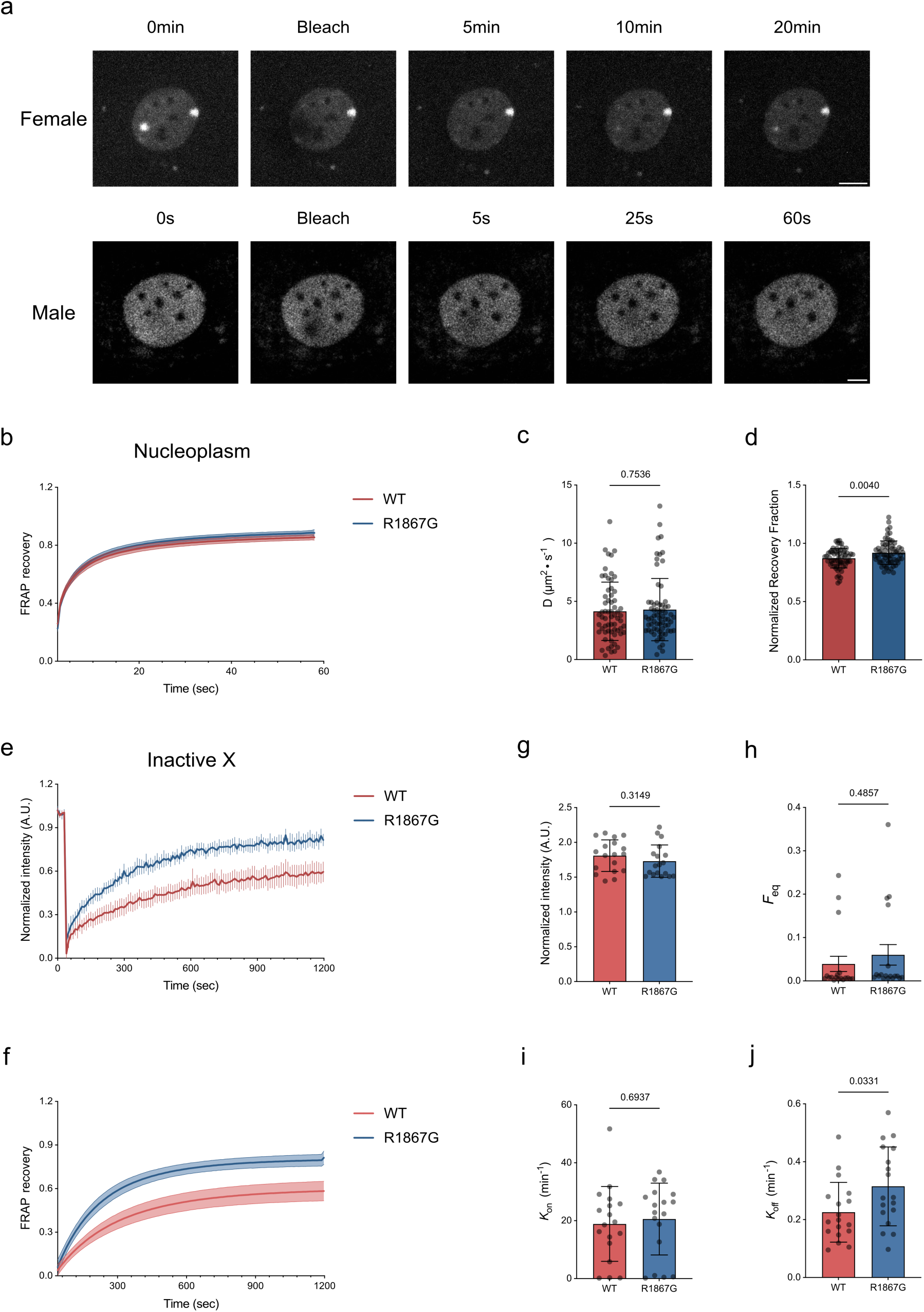
Impaired DNA binding shortens SMCHD1 chromatin residence at the Xi. a, Representative images of FRAP experiments at the Xi in female MEFs and in the nucleoplasm of male MEFs expressing SMCHD1-GFP. Scale bars, 10 µm (female cell) and 5 µm (male cell). b, Fitted nucleoplasmic FRAP recovery curves in male MEFs. Data are mean with 95% confidence intervals. c, Diffusion coefficients of WT and R1867G SMCHD1-GFP derived from model fitting. d, Normalized recovery fraction of SMCHD1-GFP within the bleached nucleoplasmic region 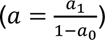. For b-d, data were pooled from three independent experiments; n = 63 WT and 67 R1867G cells. In c,d, data are mean ± s.d. and P values were calculated using two-sided t-tests. e, Normalized Xi FRAP recovery curves in female MEFs selected for comparable pre-bleach Xi fluorescence. f, Fitted Xi FRAP recovery curves for the cells shown in e. In e,f, data are mean with 95% confidence intervals. g, Normalized pre-bleach Xi fluorescence in the selected cells. h, Freely diffusing fraction of SMCHD1 at the Xi. i, Apparent association rate of freely diffusing SMCHD1 with chromatin. j, Dissociation rate of chromatin-bound SMCHD1. For e-j, data were pooled from three independent experiments; n = 18 WT and 18 R1867G cells. Data in g,i,j are mean ± s.d.; data in h are mean ± s.e.m. P values were calculated using two-sided t-tests.

Fitting the nucleoplasmic recovery curves with a circular-spot diffusion model^46, 47^ (Fig. 4b), yielded comparable diffusion coefficients for WT and R1867G SMCHD1 (Fig. 4c).

However, R1867G cells showed a larger recovery amplitude (*a*_1_) and a smaller immediate post-bleach residual fluorescence (*a*_0_) than WT cells (Extended Data Fig. 4c, d), indicating greater fluorescence recovery but also possibly a greater effective bleach depth. To account for this difference, we normalised the recovery amplitude to bleach depth. After this correction, R1867G still showed greater recovery than WT (Fig. 4d), despite having a comparable diffusion coefficient. These results indicate that a larger fraction of R1867G SMCHD1 molecules is exchangeable within nucleoplasmic regions.

We next performed FRAP at the Xi in female MEFs. The slow recovery in this compartment suggested that association and dissociation, rather than free diffusion alone, dominated the measured dynamics (Fig. 4a and Supplementary Video 2). We normalised Xi fluorescence and fitted the recovery curves using a one-binding-state model^46, 47^ (Extended Data Fig. 4f, g). In the full cell population, WT and R1867G cells showed similarly small freely diffusing fractions at the Xi (*F*_eq_; Extended Data Fig. 4i). R1867G cells displayed higher apparent association (*k*^∗^) and dissociation (*k*_off_) rates than WT cells (Extended Data Fig. 4j, k). However, the apparent association rate depends on both the concentration of unoccupied binding sites and the local concentration of fluorescent molecules^48^. Because R1867G reduced SMCHD1 abundance at the Xi in both fixed and live cells (Extended Data Fig. 4h), unequal pre-bleach intensity could confound comparison of *k*^∗^ . Indeed, despite lower pre-bleach fluorescence, R1867G and WT cells recovered similar absolute fluorescence after bleaching (Extended Data Fig. 4e).

To control for this difference, we selected WT and R1867G cells with comparable pre-bleach Xi fluorescence (Fig. 4g). R1867G cells still recovered more rapidly than WT cells (Fig. 4e, f). Model fitting showed comparable freely diffusing fractions and apparent association rates between the matched groups (Fig. 4h, i), whereas the dissociation rate remained increased in R1867G cells (Fig. 4j). The higher *k*_off_ indicates a shorter residence time for R1867G SMCHD1 at the Xi. Within the limits of the FRAP model, these results support a role for hinge-domain DNA binding in stabilizing chromatin-bound SMCHD1 after association, rather than in determining its initial recruitment to the Xi.

### DNA binding constrains local and long-range SMCHD1 mobility at the Xi

FRAP reports the ensemble dynamics of molecules within a bleached region. To resolve SMCHD1 dynamics with higher spatiotemporal resolution and to measure movement within and between the Xi and adjacent nucleoplasm, we next performed line-scan fluorescence fluctuation spectroscopy (FFS) in female MEFs (Fig. 5a). This involved applying autocorrelation function (ACF) and pair correlation function (pCF) analysis to line scan data acquired rapidly across the Xi territory^49–51^ (Fig. 5a). During rapid line scanning, the confocal detection volume moves across the selected line, encoding temporal fluorescence intensity fluctuations as SMCHD1-GFP molecules move through the detection volume at each position, which are then spatially correlated across defined distance to measure molecular movement (Fig. 5a). We calibrated the three-dimensional detection volume under our experiment conditions using fluorescein and incorporating with Gaussian point-spread function^52^ (PSF; Extended Data Fig. 5a), yielded an estimated lateral radius of approximately 200 nm and an axial radius of approximately 600 nm (Extended Data Fig. 5b).

**Fig. 5.**
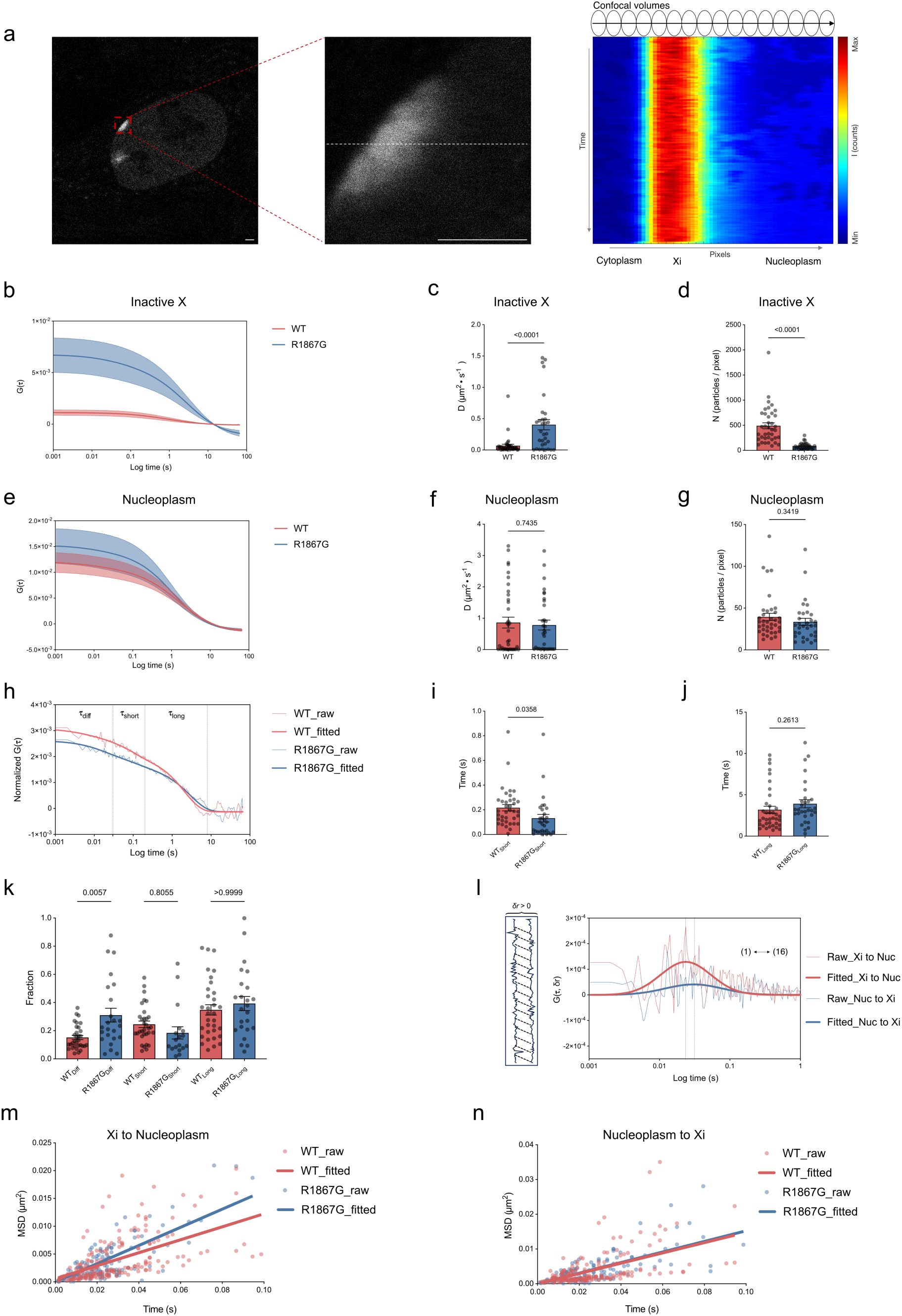
Impaired DNA binding increases local SMCHD1 mobility at the Xi and accelerates movement out of the Xi territory. a, Line-scan FFS strategy. Left, representative female MEF selected for analysis. Middle, enlarged Xi territory; the dashed white line indicates the scan path and direction. Right, carpet plot of detrended fluorescence fluctuations along the scanned line. Ovals denote confocal detection volumes, and the arrow indicates the scan direction. Scale bar, 2 µm. b, Fitted autocorrelation-function (ACF) curves within the Xi. Data are mean with 95% confidence intervals. c, Apparent SMCHD1 diffusion coefficient within the Xi. d, Estimated SMCHD1 abundance within the Xi detection volume. e, Fitted ACF curves in the nucleoplasm. Data are mean with 95% confidence intervals. f, Apparent SMCHD1 diffusion coefficient in the nucleoplasm. g, Estimated SMCHD1 abundance within the nucleoplasmic detection volume. For b-g, n = 38 WT and 31 R1867G cells. Data in c,d,f,g are mean ± s.e.m.; P values were calculated using two-sided t-tests. h, Representative ACF fit using a model incorporating freely diffusing, short-lived bound and long-lived bound states. *τ*diff, three-dimensional diffusion time; *τ*short, residence time of the short-lived bound state; *τ*long, residence time of the long-lived bound state. i,j, Residence time of the short-lived (i) and long-lived (j) SMCHD1 populations at the Xi. Data are mean ± s.e.m.; n = 37 WT and 30 R1867G cells. P values were calculated using two-sided t-tests. k, Fractions of freely diffusing, short-lived bound and long-lived bound SMCHD1 at the Xi. Data are mean ± s.e.m. P values for the indicated comparisons were calculated by one-way ANOVA with Bonferroni correction. l, Representative pCF curve fitted with a Gaussian probability distribution. Dashed lines indicate the correlation peak, and the corresponding lag time represents the mean arrival time between the two positions. An offset of 16 pixels (approximately 1,172 nm) is shown. m, Mean squared displacement analysis of SMCHD1 movement from the Xi to the nucleoplasm. Directional diffusion coefficients: WT, 0.0292 µm²·s⁻¹; R1867G, 0.0409 µm²·s⁻¹. n, Mean squared displacement analysis of SMCHD1 movement from the nucleoplasm towards the Xi. Directional diffusion coefficients: WT, 0.037 µm²·s⁻¹; R1867G, 0.038 µm²·s⁻¹.

We first analysed local SMCHD1 dynamics using ACF analysis of the line-scan data, which correlated fluorescence intensity fluctuations at the same spatial location over time. Five consecutive pixels within the Xi region were selected, in this environment that is visible from the fluorescence-intensity carpet. Then calculating their averaged ACF and finally fitting the recovered average ACF profile with a 3D diffusion model, with the decay and amplitude of the ACF providing estimates of SMCHD1’s local diffusion coefficient and concentration, respectively^53^ (Fig. 5b and Extended Data Fig. 5c). This analysis revealed WT SMCHD1 to have a slow local diffusion rate while R1867G mutant has a substantially higher apparent diffusion coefficient within the Xi (WT, 0.06843 µm²·s⁻¹; R1867G, 0.4046 µm²·s⁻¹; Fig. 5c), indicating increased local mobility of R1867G mutant. Estimated molecular abundance within the Xi detection volume was also reduced in R1867G cells (Fig. 5d), consistent with the imaging and ChIP-seq results. We then performed the same ACF analysis in nucleoplasmic regions adjacent to the Xi following the same protocol described for within the Xi (Fig. 5e and Extended Data Fig. 5d) and found both proteins diffused faster in the nucleoplasm than within the Xi, and their nucleoplasmic diffusion coefficients were comparable (WT, 0.8616 µm²·s⁻¹; R1867G, 0.7831 µm²·s⁻¹; Fig. 5f), consistent with the nucleoplasmic FRAP analysis (Fig. 4b, c). The estimated SMCHD1 abundance was markedly lower in the nucleoplasm than at the Xi and did not differ between WT and R1867G cells (Fig. 5g). The selective increase in R1867G local mobility within the Xi is therefore consistent with DNA binding constraining SMCHD1 specifically in regions containing abundant SMCHD1 binding sites.

The high density of SMCHD1 at the Xi suggested that the population may comprise multiple dynamic states, as reported for RNA polymerase II within the Xi territory^54^. We therefore fitted the ACF profiles derived from the Xi territory with a modified diffusion model that incorporated short-lived bound and long-lived bound states^55, 56^ (Fig. 5h). The fractions assigned to the short- and long-lived bound states did not differ significantly between WT and R1867G cells, whereas the freely diffusing fraction was increased in R1867G cells (Fig. 5k). R1867G also shortened the residence time of the short-lived bound population, while leaving the residence time of the long-lived population unchanged (Fig. 5i, j). Thus, impaired DNA binding shifts a subset of Xi-associated SMCHD1 from a transiently bound state towards a more mobile state, further supporting a role for DNA interactions in stabilizing SMCHD1 on the Xi chromatin.

We next tested whether DNA binding influences SMCHD1 exchange between the Xi and the surrounding nucleoplasm using pCF analysis that correlates SMCHD1 dynamics adjacent to the Xi with SMCHD1 dynamics within the Xi in a direction-dependent manner^53^. To do so, we selected one position within the Xi and a second position in the nucleoplasm at a defined spatial offset (*δ*_r_) and calculated the pCF to enter versus exit this environment. Fitting the resulting pCF curve with a Gaussian probability distribution yielded an arrival time (*τ*), representing the mean transit time required for correlated SMCHD1 fluctuations to traverse *δ*_r_ (Fig. 5l). For spatially resolved pCF analysis, *δ*_r_ must exceed the lateral radius of the confocal detection volume, such that two observation volumes are spatially independent. The line-scan pixel size was 73.22 nm in our line-scan FFS experiments. Therefore, offsets of four pixels or more exceeded the calibrated lateral radius of 213.7 nm (Extended Data Fig. 5b). Analysis at individual offsets of 8 (584 nm), 10 (732 nm), 12 (876 nm) and 16 (1168 nm) pixels detected no significant genotype-dependent difference in arrival time in either the Xi-to-nucleoplasm or nucleoplasm-to-Xi direction (Extended Data Fig. 5e-l).

To capture transport behaviour across a broader spatial range, we extended the analysis to a variable offset of 4, 8, 12, 16, 20, 24, 28 and 32 pixels (292, 584, 876, 1168, 1460, 1752, 2044, 2336 nm) and plotted mean squared displacement (MSD) as a function of arrival time^8, 53^. The approximately linear relationships in both directions were consistent with a freely diffusing component moving between the Xi and the nucleoplasm, despite the large immobile population within the Xi (Fig. 5m, n). In the Xi-to-nucleoplasm direction, R1867G SMCHD1 showed a higher directional diffusion coefficient than WT SMCHD1 (WT, 0.0292 µm²·s⁻¹; R1867G, 0.0409 µm²·s⁻¹; Fig. 5m), indicating that DNA binding slows SMCHD1 unloading from the Xi. By contrast, directional diffusion from the nucleoplasm towards the Xi was similar for WT and R1867G SMCHD1 (WT, 0.037 µm²·s⁻¹; R1867G, 0.038 µm²·s⁻¹; Fig. 5n), providing no evidence that DNA binding accelerates SMCHD1 movement onto the Xi. R1867G also showed a modestly higher directional diffusion coefficient between two nucleoplasmic positions (WT, 0.034 µm²·s⁻¹; R1867G, 0.041 µm²·s⁻¹; Extended Data Fig. 5m), potentially reflecting interactions with autosomal chromatin within the sampled region. Together, the ACF and pCF analyses show that DNA binding restrains both SMCHD1 local mobility within the Xi and long-range movement out of the Xi territory, while having little detectable effect on SMCHD1 movement towards the Xi.

### DNA binding stabilizes SMCHD1 at the Xi during its mitotic dissociation and post-mitotic accumulation

Our FRAP and FFS experiments characterized SMCHD1 dynamics during interphase. We next examined how SMCHD1 associates with the Xi during mitosis. To track dividing cells, we performed long-term lattice light-sheet imaging of live female NSCs, which divide more rapidly than primary MEFs^57, 58^. SMCHD1-GFP was lost from the nucleus before cell division and re-accumulated in the daughter-cell nuclei after mitosis (Extended Data Fig. 6a and Supplementary Video 3). At metaphase, SMCHD1-GFP was dispersed throughout the cytoplasm as the nuclear envelope broke down and chromosomes adopted a compact, rod-like morphology^59, 60^ (Extended Data Fig. 6a), indicating that SMCHD1 was not detectably retained on mitotic chromatin. We aligned individual time courses to the nuclear envelope breakdown timepoint (NEBD, t = 0). WT and R1867G cells both showed a sharp decrease in nuclear SMCHD1-GFP at nuclear envelope breakdown timepoint, followed by gradual recovery during mitotic exit and early interphase (Extended Data Fig. 6c). We used the live-cell DNA dye SPY650-DNA to visualize mitotic DNA dynamics. Mean DNA intensity increased during chromosome condensation and decreased after formation of the daughter nuclei (Extended Data Fig. 6d), consistent with changes in DNA compaction during mitosis^61^.

SMCHD1 was also lost from the Xi, but its Xi-associated signal declined gradually before the abrupt loss of bulk nuclear fluorescence at the time of nuclear envelope breakdown (Fig. 6a, b). The Xi forms a compact heteropyknotic Barr body during interphase and undergoes further chromosome condensation during mitosis^62^, which was evident in the live-cell images (Fig. 6a). To test whether Xi compaction preceded SMCHD1 loss, we quantified the coefficient of variation of DNA fluorescence within the Xi territory as a measure of the Xi chromosome heterogeneity and condensation (Extended Data Fig. 6e). Xi-associated SMCHD1 began to decline approximately 40 min before nuclear envelope breakdown, when DNA remained relatively homogeneous within the SMCHD1-defined Xi territory. Detectable DNA condensation occurred later, approximately 10-20 min before nuclear envelope breakdown, in both WT and R1867G cells (Fig. 6a, b and Extended Data Fig. 6e). Thus, the initial loss of SMCHD1 from the Xi precedes pronounced Xi condensation and is unlikely to be triggered solely by physical chromosome compaction. Re-accumulation at the Xi after division was slower than dissociation before mitosis (Fig. 6b). Nevertheless, SMCHD1 could be detected at the Xi during cytokinesis, before bulk cytoplasmic SMCHD1 had fully re-entered the daughter nuclei (Extended Data Fig. 6b), indicating that SMCHD1 Xi recruitment begins early during mitotic exit.

**Fig. 6.**
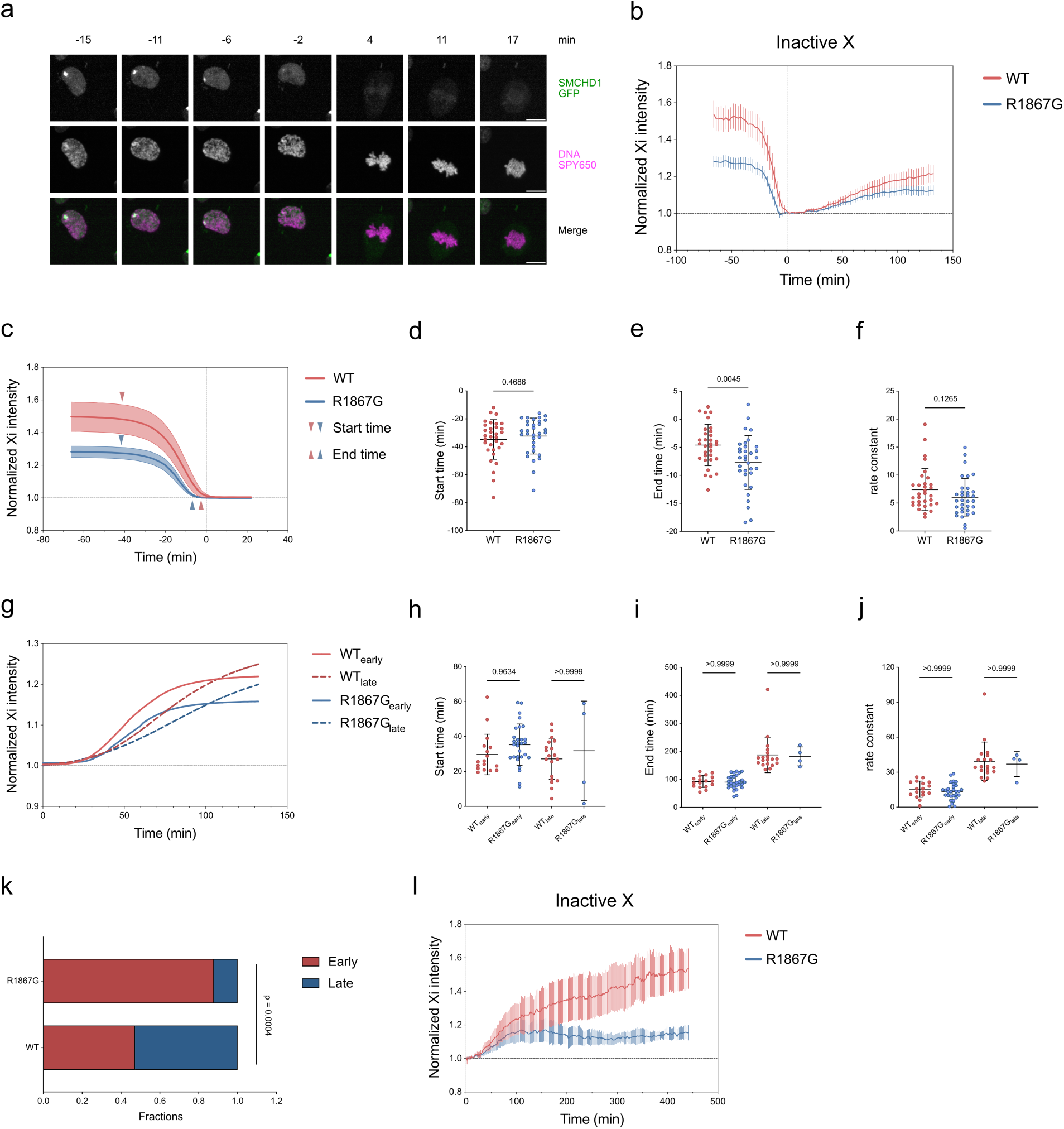
DNA binding stabilizes SMCHD1 release from the Xi during mitosis and ensures its long-term accumulation afterwards. a, Representative live-cell images showing gradual loss of SMCHD1-GFP from the Xi before division in female NSCs. Scale bar, 10 µm. b, Normalized Xi SMCHD1-GFP fluorescence before and after nuclear-envelope breakdown (t = 0). Mean Xi fluorescence was normalized to mean nuclear fluorescence (*I*xi / *I*Total Nucleus). Data are mean with 95% confidence intervals; n = 48 WT and 52 R1867G cells. c, Fitted curves for the descending phase in b. The start and end times denote the points at which each curve reached 95% and 5% of its fitted dynamic range, respectively. Data are mean with 95% confidence intervals. d-f, Fitted dissociation start time (d), end time (e) and rate constant (f). Data are mean ± s.d.; P values were calculated using two-sided t-tests. g, Fitted curves for the ascending phase in b. Cells were classified as early- or late-plateau according to whether the fitted upper plateau was reached within the observation period. For clarity, group means are shown. h-j, Fitted accumulation start time (h), end time (i) and rate constant (j). Data are mean ± s.d.; P values for the indicated comparisons were calculated by one-way ANOVA with Bonferroni correction. k, Proportion of WT and R1867G cells assigned to the early-and late-plateau populations. n = 36 WT and 33 R1867G cells; the P value was calculated using Fisher’s exact test. l, Normalized Xi SMCHD1-GFP fluorescence during long-term imaging after cell division. Data are mean with 95% confidence intervals; n = 18 WT and 17 R1867G cells. Data were pooled from two independent experiments.

To determine whether DNA binding influences SMCHD1 dissociation and re-accumulation at the Xi, we fitted the descending and ascending phases of the normalized Xi fluorescence trajectories separately using Gompertz sigmoidal models^63^. During dissociation, R1867G cells had a lower fitted upper plateau than WT cells (Extended Data Fig. 6g), consistent with reduced interphase Xi enrichment. We defined the onset and completion of dissociation as the times at which the fitted curve crossed 95% and 5% of its dynamic range, respectively. WT and R1867G cells initiated SMCHD1 loss at similar times (Fig. 6d), but R1867G cells reached the lower plateau earlier (Fig. 6e). Most cells completed SMCHD1 loss before NEBD, although a subset retained detectable Xi-associated SMCHD1 after NEBD (Extended Data Fig. 6j), indicating that Xi dissociation is temporally associated with, but not strictly determined by, nuclear-envelope breakdown. Across all cells, the fitted dissociation rate constant did not differ between genotypes (Fig. 6f). Because sigmoidal rate estimates can depend on plateau amplitude^64^, we repeated the analysis in WT and R1867G cells with comparable pre-mitotic Xi fluorescence (Extended Data Fig. 6f,h). In this matched subset, R1867G cells lost SMCHD1 from the Xi more rapidly than WT cells (Extended Data Fig. 6i). These data indicate that DNA binding delays SMCHD1 dissociation from the Xi as cells enter mitosis.

We next analysed SMCHD1 re-accumulation at the Xi in daughter cells. R1867G trajectories more often appeared to reach a plateau within the 2-h observation window, whereas many WT trajectories continued to increase (Fig. 6b and Extended Data Fig. 6k). Individual cells were separated into an early-plateau population, which reached the fitted upper plateau during the observation period, and a late-plateau population, for which the fitted plateau was reached only after the observation period (Fig. 6g). Within both populations, WT and R1867G cells initiated SMCHD1 Xi accumulation at similar times after division (Fig. 6h), reached their respective plateaus at similar times (Fig. 6i), and displayed comparable accumulation-rate constants (Fig. 6j). Thus, impaired DNA binding did not measurably affect the onset or kinetics of initial SMCHD1 recruitment to the Xi. However, a significantly greater fraction of R1867G cells belonged to the early-plateau population (Fig. 6k), indicating that rather than affecting initial recruitment, impaired DNA binding affects high level SMCHD1 Xi maintenance after mitosis. Consistent with this, long-term imaging for approximately 7 h after cell division showed continued accumulation of WT SMCHD1 at the Xi, whereas R1867G reached an earlier plateau and failed to sustain high-level Xi enrichment (Fig. 6l). Together, these observations show that DNA binding is dispensable for the initial recruitment of SMCHD1 to newly formed Xi territories but is required to sustain its progressive accumulation and long-term retention during interphase.

## Discussion

Here, we have used live cell imaging and genomic analyses to study SMCHD1 function, specifically how the hinge domain DNA binding activity contributes to SMCHD1’s function on chromatin. Our results support a model in which DNA binding does not primarily drive SMCHD1 recruitment, but instead stabilizes chromatin-bound SMCHD1, restricts its mobility and enables full SMCHD1 function in cells (Fig. 7).

**Fig. 7.**
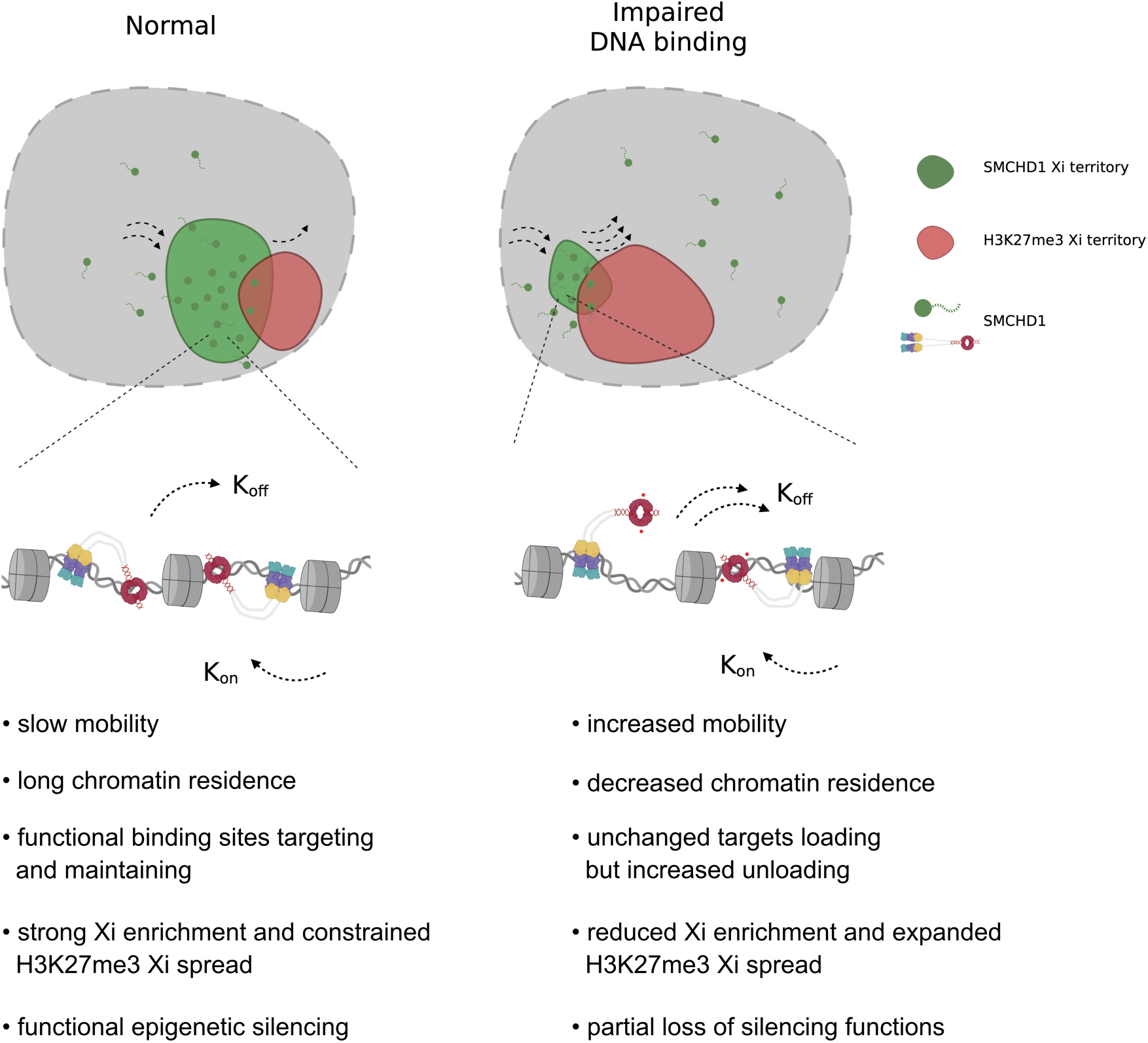
Hinge domain DNA binding enables SMCHD1 chromatin residence and functions. Model showing that SMCHD1 is initially recruited to chromatin through mechanisms that remain incompletely defined, potentially involving chromatin marks, interacting proteins and ATPase-dependent processes. Hinge-domain DNA binding does not appear to act as the primary recruitment mechanism. Instead, DNA binding stabilizes chromatin-associated SMCHD1, constrains its mobility and prolongs its residence at binding sites. This maintenance function supports robust SMCHD1 accumulation at the inactive X chromosome and contributes to proper regulation of Xi chromatin state. When DNA binding is impaired by the R1867G mutation, SMCHD1 remains capable of chromatin association but becomes less stably retained and more mobile, leading to reduced Xi enrichment, increased movement out of the Xi territory and expanded H3K27me3 spreading across the Xi. Thus, DNA binding maintains SMCHD1 on chromatin and enables full SMCHD1 function.

Impaired DNA binding reduced SMCHD1 enrichment at the Xi territory in both female MEFs and NSCs, without impacting overall amounts of SMCHD1. In contrast, SMCHD1 occupancy at many autosomal sites was only modestly affected, indicating that hinge-domain mutant SMCHD1 retains the capacity to associate with chromatin and occupy a subset of SMCHD1-bound regions. Consistent with this, FLIM-FRET analysis showed that WT and R1867G SMCHD1 exhibited comparable transient interaction frequencies with H2B across the nucleus in living male cells. Analogous to the multiple DNA-binding modes described for transcription factors, which can include both non-specific and sequence-specific interactions^55, 65, 66^, we propose that SMCHD1 may engage chromatin through a spectrum of binding states, despite lacking strong sequence-specific DNA-binding preference^16^. Impaired DNA binding shifts SMCHD1 towards a less stable, more diffusive state at the Xi, whereas binding at some autosomal regions may be maintained by additional mechanisms, including interactions with other proteins, chromatin features or local chromatin compaction. A limitation of our stringent ChIP-seq approach is that it preferentially detects stable or highly enriched binding events and may not capture short-lived or transient SMCHD1-chromatin interactions. Higher-sensitivity or kinetic chromatin-profiling approaches will be needed to define these transient binding states genome-wide in future.

Complementary live-cell imaging approaches revealed how DNA binding affects SMCHD1 dynamics. In nucleoplasmic regions of male cells, impaired DNA binding did not measurably alter the SMCHD1 diffusion coefficient, but increased fluorescence recovery after photobleaching, consistent with reduced stable chromatin association. At the Xi territories, R1867G showed a comparable association rate but a faster dissociation rate from chromatin-bound sites, indicating that DNA binding prolongs SMCHD1 residence rather than mediating initial recruitment. FFS analysis further showed reduced binding time, an increased fraction of diffusive molecules and a higher local diffusion rate for R1867G within the Xi territories. Importantly, pCF analysis showed that R1867G diffused out of the Xi more rapidly than WT SMCHD1, whereas movement from nucleoplasm into the Xi was comparable. Together, these findings indicate that DNA binding maintains SMCHD1 association with chromatin rather than recruiting SMCHD1, acting at both molecular binding-site and chromosome-territory scales.

The extensive enrichment of SMCHD1 on the Xi is unlikely to arise instantaneously but instead develops progressively. SMCHD1 is not enriched on the Xi in undifferentiated cells, but accumulates gradually during differentiation, indicating that progression of X chromosome inactivation and establishment of the Xi chromatin environment are important for SMCHD1 recruitment and accumulation^30, 67^. H2AK119ub, deposited through the Xist-HNRNPK-PRC1 pathway, is a central upstream determinant of SMCHD1 enrichment at the Xi, both during the establishment of X inactivation during differentiation and in the maintenance phase post-differentiation^24, 26^. SMCHD1-interacting proteins may also contribute to its Xi enrichment. One candidate is LRIF1, which is required for SMCHD1 enrichment at pericentric heterochromatin by bridging SMCHD1 and HP1^68^. Loss of LRIF1 in undifferentiated cells prevents subsequent SMCHD1 enrichment at the Xi after differentiation, whereas LRIF1 deletion in differentiated cells has little effect^30, 68^. These data suggest LRIF1 also contributes to SMCHD1 recruitment during establishment of X inactivation but no longer plays a role during the maintenance phase. From a protein-dynamics perspective, SMCHD1 accumulation on the Xi reflects both recruitment to the Xi territory and maintenance at chromatin-bound sites. Defects in either process could reduce SMCHD1 Xi enrichment. Consistent with this, Constantinescu et al. recently reported that impaired ATP hydrolysis reduces SMCHD1 enrichment on the Xi and shortens chromatin residence time, suggesting that ATPase activity contributes to SMCHD1 maintenance on the Xi^30^. However, whether ATPase activity also contributes to the recruitment process, and whether altered Xi chromatin state contributes to reduced SMCHD1 enrichment, both remain unclear. In our study, the switchable system allowed endogenous SMCHD1 to be replaced with mutant SMCHD1 after X chromosome inactivation had already been established. We cannot exclude the possibility that secondary changes in the Xi chromatin state contributed to the phenotypes observed in this system. However, such effects are likely to be less pronounced than those caused by perturbing SMCHD1 before or during X chromosome inactivation. Consistent with this, we did not observe reactivation of X-linked genes in our system or in comparable post-establishment settings^17, 24^. We therefore favour a model in which the reduced Xi enrichment of R1867G is caused primarily by impaired DNA binding, which compromises maintenance of chromatin-bound SMCHD1 rather than its initial recruitment at the molecular level.

To more fully explore recruitment versus maintenance of SMCHD1 on the Xi we also characterized SMCHD1 dynamics during mitosis. We found that SMCHD1 dissociates from mitotic chromosomes, including the inactive X chromosome. Temporal kinetic analysis of SMCHD1 loss from the Xi showed that R1867G enrichment declined more rapidly than WT SMCHD1, indicating that DNA binding contributes to SMCHD1 maintenance at both interphase and the pre-mitotic Xi. We further observed gradual re-accumulation of SMCHD1 at the Xi in daughter cells after division. Importantly, impaired DNA binding did not affect the initial accumulation of SMCHD1 on the Xi, supporting our conclusion that DNA binding is not required for the recruitment step. Instead, R1867G failed to sustain continued SMCHD1 accumulation over longer post-mitotic periods, again highlighting the role of DNA binding in SMCHD1 retention on the chromatin. The biological significance of mitotic SMCHD1 unloading remains to be determined, but it may be linked to proper mitotic chromosome regulation. During mitosis, *Xist* and several *Xist*-associated proteins dissociate from the inactive X chromosome, whereas high levels of H3K27me3 are retained^69–71^. This mitotic Xi identity is maintained in part through differential acetyltransferase occupancy and acetylation levels between Xi and Xa^71^. It is therefore possible that unloading of SMCHD1 from the Xi is required for proper establishment or maintenance of the mitotic Xi H3K27me3 profile, thereby helping to preserve the Xi identity during mitosis. How SMCHD1 is rapidly unloaded from chromosomes during mitosis also remains to be explored in future work. Our data suggest that this process is unlikely to be driven simply by physical chromosome condensation, at least at the inactive X chromosome. Inhibition of Aurora kinase B has been shown to impair mitotic *Xist* release without altering histone H3 phosphorylation^69^. The *Xist*-associated protein CIZ1 also disassembles from the Xi during mitosis through an Aurora kinase B-regulated, phosphorylation-dependent mechanism^72^. Whether SMCHD1 unloading is controlled by mitotic kinase signalling remains an important question for future studies.

## Methods

### Generation of LSL-SMCHD1-GFP knock-in mouse strains

A targeting plasmid was generated containing full-length mouse *Smchd1* cDNA, either wild type or harbouring the R1867G mutation, with AcGFP fused to the C terminus through a GSSG linker. A loxP-stop-loxP (LSL) cassette was inserted 5′ of the *Smchd1* cDNA to enable Cre-dependent expression. The expression cassette was flanked by homology arms targeting the *Rosa26* locus. Knock-in mouse strains were generated by the MAGEC mouse genetic engineering facility at the Walter and Eliza Hall Institute of Medical Research (WEHI). Cas9 protein, a *Rosa26*-targeting sgRNA and the targeting plasmid used as a repair template were microinjected into single-cell fertilized embryos. The sgRNA sequence CTCCAGTCTTTCTAGAAGAT was used to introduce double-strand breaks at the *Rosa26* locus and stimulate homology-directed repair. Founder mice were generated on a C57BL/6 background and bred to wild-type C57BL/6 mice for more than five generations while selecting for the knock-in allele, to minimize the retention of potential off-target variants introduced during genome editing. Correct integration of the LSL-SMCHD1-GFP knock-in cassettes (both wild-type and R1867G mutant) were confirmed using both long-range and short-range PCR primer sets, followed by Illumina sequencing. The LSL-SMCHD1-GFP knock-in mouse strains were crossed with *Smchd1* conditional knockout mice^17, 73^ (*Smchd1*^fl/fl^), along with the *Rosa:CreERT2* knockin^39^ to generate Cre-inducible switchable SMCHD1 strains. Animals were genotyped by PCR with primers provided in Supplementary Table 1.

All lines were bred on the C57BL/6 inbred background. All animal procedures were performed in accordance with institutional guidelines and were approved by the WEHI Animal Ethics Committee, under approval numbers AEC 2020.050, 2020.048, 2023.033 and 2023.045.

### Derivation of MEF and NSC lines

Mouse embryos were collected at embryonic day 11.5-14.5 for derivation of MEFs. The uterus was removed, and individual embryos were dissected and washed in cold PBS.

The embryo head and internal organs were removed, and one forelimb was collected for genotyping. The remaining embryo body was mechanically dissociated by vigorous pipetting in DMEM supplemented with 10% (v/v) fetal bovine serum (FBS) to generate a single-cell suspension. Cells were cultured in DMEM with 10% (v/v) FBS.

For NSC derivation, embryos were collected at embryonic day 14.5. Embryo heads were isolated, and the meninges and subcortical tissue were removed. Forebrain cortical regions were collected and mechanically minced. Cortical tissue pieces were transferred into NeuroCult^TM^ NSC Basal Medium (STEMCELL Technologies) containing 1 mg/ml DNase I and 0.5% trypsin–EDTA, and incubated at 37 °C with 5% CO_2_ for 20 min. Following digestion, cortices were mechanically dissociated by repeated pipetting until a single-cell suspension was obtained.

### Cell culture and treatment

Primary MEFs were cultured under low-oxygen conditions in DMEM supplemented with 10% (v/v) FBS at 37 °C, 5% O_2_ and 5% CO_2_. Cells were detached with trypsin for passaging.

NSCs were cultured in NeuroCult^TM^ NSC Basal Medium supplemented with NeuroCult^TM^ Proliferation Supplement (STEMCELL Technologies), 20 ng/ml recombinant human EGF and 20 ng/ml recombinant human basic FGF. Laminin was added to the medium at 10 ng/ml to promote NSC adherence. NSCs were maintained at 37 °C with 5% CO_2_ and detached with Accutase for passaging. To switch expression from endogenous SMCHD1 to SMCHD1-GFP, MEFs and NSCs were treated with 1 µM 4-hydroxytamoxifen (4-OHT). Medium containing 4-OHT was replaced every 24 h with fresh 4-OHT-containing medium.

Drosophila S2 cells were purchased from Thermo Fisher Scientific (R69007) and cultured according to the manufacturer’s instructions. Briefly, S2 cells were maintained in Schneider’s medium (Thermo Fisher Scientific, 21720024) supplemented with 10% fetal calf serum (FCS) at 28 °C.

### Flow Cytometry

Neural stem cells were dissociated using Accutase and centrifuged at 1,000 rpm for 5 min at room temperature. The resulting cell pellet was resuspended in 1× KnockOut™ Serum Replacement (Life Technologies) and passed through a cell strainer into 5 mL glass tubes. Fluorescence-activated cell sorting of GFP-positive cells was carried out using a BD FACSAria™ Fusion (BD Biosciences). GFP-negative gates were defined manually using GFP-negative cells as controls. Data were analyzed using FlowJo v10 (Tree Star).

### Western blotting

For non-histone western blotting, whole-cell protein extracts were prepared as previously described^26^. Protein concentrations were determined using a standard bicinchoninic acid (BCA) assay. Total protein lysate, 25-40 µg, was mixed with 1× reducing sample buffer, boiled at 95 °C for 10 min and centrifuged at maximum speed for 1 min. Proteins were resolved using NuPAGE 3-8% Tris-acetate gels for separation of SMCHD1 and SMCHD1-GFP, or NuPAGE 4-12% Bis-Tris gels for other targets. Proteins were transferred onto PVDF membranes by wet transfer at 100 V and 4 °C for 2 h for SMCHD1 or 1 h for other targets. Membranes were blocked for 1 h at room temperature in blocking buffer comprising 5% (w/v) skim milk in PBS containing 0.1% Tween-20 (PBST). Membranes were then incubated with primary antibodies (Reporting Summary) at optimized dilutions overnight at 4 °C. After primary antibody incubation (see Supplementary Table 2 for all antibody details), membranes were washed four times for 10 min each in PBST. HRP-conjugated secondary antibodies diluted in blocking buffer were applied for 1 h at room temperature, followed by four additional washes in PBST. Signals were detected using enhanced chemiluminescence (ECL) substrate and visualized using a Bio-Rad Gel Doc imaging system.

For histone western blotting, cells were harvested and washed twice with cold PBS. Cells were lysed in Triton extraction buffer (TEB; PBS containing 0.5% Triton X-100 (v/v), 2 mM PMSF and 0.02% NaN_3_ (w/v)) on ice for 10 min with gentle agitation. Samples were centrifuged at 6,500 × g for 10 min, and the supernatant was discarded. Pellets were washed with half the original volume of TEB and centrifuged again. Nuclear pellets were resuspended in 0.2 N HCl and incubated overnight at 4 °C. Samples were then centrifuged at 6,500 × g for 10 min, and the supernatant containing acid-extracted histones was collected. Protein concentration measurement and subsequent western blotting procedures were performed as described above for non-histone samples.

### Immunofluorescence

Cells were seeded onto 13-mm glass coverslips at an appropriate density one day before fixation. Cells were washed with PBS and fixed with 4% paraformaldehyde (PFA) for exactly 10 min at room temperature. After fixation, cells were washed three times with cold PBS for 5 min each and permeabilized with 0.5% Triton X-100 on ice for 5 min. Cells were then washed three times with cold PBS and blocked in 1% BSA for 30 min at room temperature. Primary antibodies (Reporting Summary) diluted in blocking buffer were applied overnight at 4 °C in a humidified chamber. After three washes with cold PBS for 5 min each, cells were incubated with fluorophore-conjugated secondary antibodies (Reporting Summary) for 1 h at 4 °C. Cells were washed three times with cold PBS, counterstained with DAPI for 1 min at room temperature and mounted using VECTASHIELD H-1000 mounting medium (Vector Laboratories) before imaging.

Immunofluorescence imaging was performed using a Zeiss LSM 980 confocal microscope. Image acquisition was controlled using ZEN Blue 3.3 software with an auto-optimized multichannel acquisition strategy. Fixed-cell fluorescence images were acquired using a Zeiss LSM 980 laser-scanning confocal microscope equipped with a Plan-Apochromat 40×/1.3 NA oil-immersion objective. Alexa Fluor 546, Alexa Fluor 488, DAPI and Alexa Fluor 647 were excited using 561-, 488-, 405- and 639-nm laser lines, respectively. Fluorescence emission was collected over 561-641 nm for Alexa Fluor 546, 508-552 nm for Alexa Fluor 488, 408-500 nm for DAPI and 642-758 nm for Alexa Fluor 647. Alexa Fluor 546 and Alexa Fluor 488 fluorescence was detected using a spectral GaAsP-PMT detector, whereas DAPI and Alexa Fluor 647 fluorescence was detected using multialkali PMT detectors. An MBS 488/561/639 main beam splitter was used for visible excitation and an MBS -405 beam splitter for 405-nm excitation. Identical imaging settings were applied across experimental groups within each experiment. For qualitative immunofluorescence analyses, 15 optical z-sections were acquired per field. Maximum-intensity projections of z-stacks were generated for representative images. The presence of a Xi cloud was determined manually by comparing fluorescence intensity at the putative Xi region with that of the surrounding nuclear region.

For quantitative immunofluorescence analyses, images were acquired at a resolution of 1,024 × 1,024 pixels with a pixel size of 0.207 µm, two line-averaging scans and 16-bit acquisition. The number of z-sections was automatically optimized to cover the entire nuclear volume. The confocal pinhole diameter was 39.3 µm, corresponding to 1 Airy unit for the 561-nm channel. Acquired images were imported into Imaris software (Bitplane) for three-dimensional analysis. Three-dimensional surface masks of the Xi regions were generated using intensity thresholds from the SMCHD1 channel for SMCHD1 and GFP measurements, or from the H3K27me3 channel for H3K27me3 measurements. Three-dimensional nuclear surface masks were generated using the intensity threshold of the SMCHD1 channel. Quantitative parameters, including volume, mean fluorescence intensity and total fluorescence intensity, were calculated for all channels within the defined surface regions and used for downstream analyses.

### ChIP-sequencing

For SMCHD1 ChIP-seq, NSCs were treated with 4-OHT for 5 days, and GFP-positive NSCs from switchable cell lines were isolated by FACS. Sorted cells were then expanded in culture until the required cell number was obtained. For each sample, 6 × 10^7^ NSCs were harvested and washed once with cold PBS. Cells were counted and crosslinked in 40 ml fresh culture medium supplemented with 4 ml freshly prepared formaldehyde solution to achieve a final concentration of 1% formaldehyde (5 mM HEPES-KOH, pH 7.5, 10 mM NaCl, 0.1 mM EDTA and 0.05 mM EGTA). Crosslinking was performed for exactly 10 min at room temperature with gentle rotation and quenched by adding glycine to a final concentration of 125 mM. Cells were centrifuged at 500 × g for 5 min at 4 °C and washed twice with cold PBS. The supernatant was removed, and the crosslinked cell pellet was processed immediately for downstream ChIP procedures. Drosophila S2 cells were fixed using the same method, snap frozen and stored at −80 °C.

Fixed S2 cells were thawed on ice, and 1 × 10^7^ S2 cells were mixed with 6 × 10^7^ NSCs for each sample. Cells were resuspended in nuclei extraction buffer (20 mM Tris-HCl, pH 8.0, 10 mM NaCl, 2 mM EDTA and 0.5% (v/v) NP-40) supplemented with 1× cOmplete protease inhibitor cocktail (Roche) and incubated on ice for 5 min. Cells were centrifuged, the supernatant was removed, and nuclei were washed twice with nuclei extraction buffer. Isolated nuclei were resuspended in 1 ml SDS-free sonication buffer (50 mM Tris-HCl, pH 7.5, 150 mM NaCl, 5 mM EDTA, 0.5% (v/v) NP-40 and 1% (v/v) Triton X-100) supplemented with 1× cOmplete protease inhibitor cocktail. Samples were transferred into large-volume sonication tubes (Covaris milliTUBE; 3.5 × 10^7^ cells per tube) and sonicated using a Covaris ME220 sonicator with the following settings: peak power, 75 W; duty factor, 17%; cycles per burst, 1,000; duration, 1,800 s; temperature, 4-10 °C.

Sonicated samples were transferred to DNA LoBind Tube (Eppendorf, 022431021) and centrifuged at maximum speed for 20 min. The supernatant containing soluble chromatin was collected. An aliquot corresponding to 1% of the total chromatin from each sample was reserved as input. Chromatin was diluted to a final volume of 1 ml with ChIP dilution buffer (20 mM Tris-HCl, pH 8.0, 150 mM NaCl, 2 mM EDTA and 1% (v/v) Triton X-100) supplemented with 1× cOmplete protease inhibitor cocktail. Washed Protein G Dynabeads (20 µl per sample; Thermo Fisher Scientific, 10003D) were added, and samples were incubated for 2 h at 4 °C with rotation for pre-clearing. Beads and non-specifically bound material were removed using a magnetic rack. Bovine serum albumin (BSA; final concentration, 0.1%), 7.5 µg anti-GFP antibody (Thermo Fisher Scientific, A11122) and 1 µg spike-in antibody (Active Motif, 104597) were added to the pre-cleared chromatin, and samples were incubated overnight at 4 °C with gentle tumbling. The following day, 20 µl pre-washed Protein G Dynabeads were added to each sample and incubated for an additional 2 h at 4 °C with rotation. Immunocomplexes were then washed, eluted and purified as described below.

For H3K27me3 ChIP-seq, 2.5 × 10^6^ sorted GFP-positive NSCs or corresponding control NSCs were used per sample. Fixed S2 cells, 5 × 10^5^ cells per sample, were added as spike-in material. Nuclei were extracted as described above and resuspended in 130 µl ChIP buffer containing SDS (20 mM Tris-HCl, pH 7.5, 150 mM NaCl, 2 mM EDTA, 1% (v/v) NP-40 and 0.3% SDS), supplemented with 1× cOmplete protease inhibitor cocktail. Samples were transferred into Covaris sonication tubes (Covaris, 520045), and chromatin was sheared using a Covaris ME220 sonicator with the following settings: peak power, 75 W; duty factor, 27%; cycles per burst, 200; duration, 750 s; temperature, 4-10 °C. Subsequent ChIP procedures were performed as described for SMCHD1 ChIP-seq, except that 2 µg anti-H3K27me3 antibody (Cell Signaling Technology, 9733S) and 1 µg spike-in antibody (Active Motif, 104597) were used per sample.

After incubation with antibody and Protein G Dynabeads, ChIP samples were placed on a magnetic rack and the supernatant was discarded. Beads were sequentially washed once with each of the following buffers: wash buffer 1 (20 mM Tris-HCl, pH 8.0, 150 mM NaCl, 2 mM EDTA, 1% (v/v) Triton X-100 and 0.15% SDS), wash buffer 2 (20 mM Tris-HCl, pH 8.0, 500 mM NaCl, 2 mM EDTA, 1% (v/v) Triton X-100 and 0.1% SDS) and wash buffer 3 (20 mM Tris-HCl, pH 8.0, 250 mM LiCl, 2 mM EDTA, 0.5% (v/v) Igepal CA-630 and 0.5% (w/v) sodium deoxycholate). All wash buffers were supplemented with 1× cOmplete protease inhibitor cocktail. Each wash was performed for 5 min at 4 °C with gentle rotation. Beads were then washed twice briefly with TE buffer (10 mM Tris-HCl, pH 7.5 and 1 mM EDTA). Chromatin was eluted by incubating beads twice with 100 µl freshly prepared elution buffer (100 mM NaHCO_3_ and 1% SDS) for 30 min at 55-60 °C with shaking. Eluates from the same sample were combined. Input samples were adjusted to a final volume of 200 µl using elution buffer. NaCl was added to a final concentration of 200 mM, and RNase A (Sigma) was added to a final concentration of 100 µg/ml to all ChIP and input samples. Samples were incubated overnight at 65 °C with shaking to reverse crosslinks. Proteinase K (Sigma) was then added to a final concentration of 200 µg/ml, and samples were incubated for 1 h at 55 °C. DNA was purified using the Zymo ChIP DNA Clean & Concentrator kit according to the manufacturer’s instructions and stored at −20 °C until further use. Libraries were prepared using the Illumina TruSeq DNA Sample Preparation Kit according to the manufacturer’s instructions. All libraries were sequenced on an Illumina NextSeq platform to generate single-end 132-bp reads.

### ChIP-seq analysis

FASTQ files were adapter-trimmed using Trim Galore! v0.6.10 with Cutadapt v4.8, followed by quality-control assessment using FastQC v0.12.1. Trimmed reads were aligned to the mouse reference genome GRCm38/mm10 using Bowtie2 v2.5.3^74^, and alignments were processed using SAMtools v1.20^75^. PCR duplicates were removed using SAMtools and reads mapping to blacklisted genomic regions were excluded using the mm10 blacklist^76^. Reads were also independently aligned to the *Drosophila melanogaster* dm6 genome using the same pipeline to enable spike-in normalization.

BigWig files were generated from BAM files using the bamCoverage function in deepTools v3.5.5^77^. Reads were extended to 180 bp, normalized as reads per kilobase per million mapped reads (RPKM), and binned at 20-bp resolution. Mouse BigWig tracks were normalized using Drosophila S2 spike-in reads according to the following scaling factor:

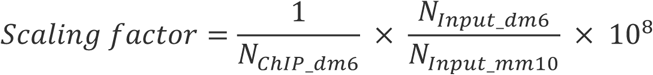

where N represents the number of uniquely mapped reads to either the mouse genome (mm10) or Drosophila genome (dm6).

Peak calling was performed using MACS3^40^ with the corresponding input sample as the control for each group and a false discovery rate (FDR) threshold of 0.05. Peaks detected in at least two biological replicates were defined as reproducible peaks. Reproducible peaks were merged using BEDTools v2.31.1^78^ and BEDOPS v2.4.41^79^. Heatmaps and average profile plots were generated using deepTools v3.5.5. Differential binding analysis was performed using the csaw package^80^ in R with an FDR threshold of 0.05.

### RNA sequencing

Control (CON; *Smchd1*^fl/fl^, *Rosa26*^+/+^), knockout (KO; *Smchd1*^fl/fl^, *Rosa26*^CreERT2/+^), WT and R1867G switchable NSC lines (*Smchd1*^fl/fl^, *Rosa26*^LSL-SMCHD1-GFP/CreERT2^) were treated with 4-OHT for 5 days. GFP-positive cells from WT and R1867G lines were isolated by FACS. For each line, 1 × 10^6^ cells were collected, and total RNA was extracted in parallel using the Zymo Quick-DNA/RNA Miniprep Plus Kit (Zymo Research, D7003). RNA quantity and integrity were assessed using NanoDrop, Qubit and TapeStation. Poly(A)^+^ mRNA enrichment and library preparation were performed using the Illumina TruSeq RNA Library Preparation Kit, non-stranded, according to the manufacturer’s instructions.

FASTQ files were adapter-trimmed using Trim Galore! v0.6.10 with Cutadapt v4.8, followed by quality-control assessment using FastQC v0.12.1. Trimmed reads were aligned to the GRCm39/mm39 mouse reference genome using HISAT2 v2.2.1^81^. BAM files were generated and processed using SAMtools v1.20. PCR duplicates were retained. Gene-level read counts were quantified in R using the featureCounts function from the Rsubread package^82^. Differential gene expression analysis was performed using edgeR^83^, with a false discovery rate (FDR) threshold of 0.05. Gene set enrichment testing was performed using the ROAST function from the limma package^84, 85^.

### Fluorescence lifetime imaging microscopy coupled with Förster resonance energy transfer (FLIM-FRET)

NSCs were treated with 4-OHT for 5 days and seeded onto 35-mm glass-bottom dishes (FluoroDish, FD35-100). Cells were transiently transfected with 1.5 µg plasmid encoding H2B-mCherry using Lipofectamine 3000 (Thermo Fisher Scientific), according to the manufacturer’s instructions. 4-OHT-treated NSCs that were not transfected with H2B-mCherry were prepared in parallel as donor-only controls. Live-cell FLIM-FRET measurements were performed 24 h after transfection.

All FLIM-FRET measurements were performed on an Olympus FV3000 confocal laser-scanning microscope equipped with a 488-nm pulsed laser operating at an 80-MHz repetition rate and coupled to an ISS A320 FastFLIM module for fluorescence lifetime acquisition. SMCHD1-GFP served as the FRET donor in all experiments and was selectively excited using the external 488-nm pulsed laser. Donor fluorescence emission was directed through a 405/488/561-nm dichroic mirror to an external photomultiplier tube detector (H7422P-40, Hamamatsu) fitted with a 500/25-nm bandpass emission filter. Before lifetime acquisition, dual-channel intensity images of SMCHD1-GFP and H2B-mCherry were acquired using identical laser, dichroic and detector settings to confirm co-expression and appropriate acceptor-to-donor ratios. FLIM data were acquired using a ×60 water-immersion objective with a numerical aperture of 1.2 at 37 °C. Regions of interest (ROIs) approximately 20-25 µm in diameter were selected within individual nuclei showing robust co-expression of donor and acceptor fluorophores. Fluorescence lifetime data were integrated over 20-30 sequential frames at a resolution of 256 × 256 pixels, with a pixel size of approximately 80 nm and a pixel dwell time of 20 µs. Instrument response calibration and lifetime fitting were performed using ISS VistaVision software, with fluorescein at pH 9, which has a single-exponential lifetime of approximately 4 ns, used as the reference standard.

Mean GFP intensity (*I_GFP_*) and mean mCherry intensity (*I_mC_*_ℎ*erry*_) were measured across the whole NSC nucleus. Cells with comparable donor-to-acceptor ratios, defined as 0.2< *I_GFP_*⁄*I_mC_*_ℎ*erry*_ <0.6, were selected for downstream analysis. Donor fluorescence lifetime was determined by phasor-based approach performed in SimFCS (Laboratory for Fluorescence Dynamics)^86, 87^. Pixel-wise FRET efficiency values were converted into chromatin-compaction maps, in which pixels with reduced donor lifetime were classified as FRET-positive and interpreted as compacted chromatin, whereas pixels lacking FRET were interpreted as more open chromatin states. Compaction maps were pseudocoloured to represent open chromatin in teal and compact chromatin in red. To analyse chromatin organization within the Xi, SMCHD1-GFP intensity images were thresholded to generate binary masks defining Xi regions. These masks were applied to the FLIM-FRET compaction maps to quantify the proportion of FRET-positive pixels within the Xi for each cell.

### Fluorescence recovery after photobleaching (FRAP)

MEFs were cultured, treated with 4-OHT for 5 days and seeded into 8-well chambered coverslips (ibidi, 80826) or 35-mm µ-Dishes (ibidi, 82166). Before imaging, culture medium was replaced with FluoroBrite^TM^ DMEM (Life Technologies, A1896701), and cells were equilibrated in a 37 °C environmental microscopy chamber for 30 min. FRAP experiments were performed using a Zeiss LSM 980 confocal microscope controlled by ZEN Blue 3.3 software.

For Xi FRAP experiments, FRAP experiments were performed on a Zeiss LSM 980 laser-scanning confocal microscope equipped with a Plan-Apochromat 40×/1.3 NA oil-immersion objective and an environmental chamber maintained at 37 °C and 5% CO_2_. SMCHD1-GFP was excited using a 488-nm laser and fluorescence was collected between 525 and 672 nm using a multialkali PMT detector. An MBS 488/639 main beam splitter was used. Images were acquired at 512 × 512 pixels with a pixel size of 0.207 µm, 2× optical zoom and a 113.8-µm pinhole corresponding to approximately 2 Airy units. Ten-plane Z stacks were acquired at 1.01-µm intervals every ∼10 s for 121 time points (∼20 min), using bidirectional scanning and twofold frame averaging. A circular region of interest approximately 2.61 µm in diameter was photobleached using the 405- and 488-nm laser lines at 100% transmission, with ten bleaching iterations and two repetitions. Fluorescence recovery was subsequently monitored under the same acquisition settings.

For nucleoplasmic FRAP experiments, male MEFs were used. Random intranuclear regions lacking visible nuclear speckles were selected for photobleaching. FRAP experiments were performed using a Zeiss LSM 980 laser-scanning confocal microscope equipped with a Plan-Apochromat 40×/1.3 NA oil-immersion objective and an environmental chamber set to 37 °C and 5% CO_2_. SMCHD1-GFP was excited using a 488-nm laser, and fluorescence was detected between 525 and 672 nm using a multialkali PMT detector. An MBS 488/639 main beam splitter was used. Single-plane images were acquired at 512 × 512 pixels with a pixel size of 0.104 µm, 4× optical zoom and a 113.8-µm pinhole corresponding to approximately 2 Airy units. Images were acquired using bidirectional scanning and twofold frame averaging at approximately 0.6-s intervals for ∼60 s. A circular region of interest approximately 2 µm in diameter was photobleached using the 405- and 488-nm laser lines at 100% power setting for 20 iterations with two repetitions, after which fluorescence recovery was monitored under the same imaging conditions.

FRAP image processing and quantitative analysis were performed using FIJI software. For Xi FRAP experiments, maximum-intensity projections were generated from z-stack images. Image sequences were registered using the StackReg and TurboReg plugins^88^. The Xi region in pre-bleach images was manually identified based on local fluorescence enrichment. Fluorescence intensity within the Xi region (*I*_Xi_) was measured over time. Fluorescence intensities from an identically sized nuclear region (*I*_nuc_) and from a region outside the nucleus (*I*_back_) were also measured to correct for whole-nucleus fluorescence changes and background signal, respectively. Cells with excessive movement that could not be reliably corrected by image registration were analysed using TrackMate 7^89^. In these cases, Xi and nuclear regions were segmented by intensity thresholding and tracked over time to extract fluorescence intensity values. Background fluorescence was measured from a defined region outside the nucleus.

Xi fluorescence intensities were normalized using a double-normalization approach:

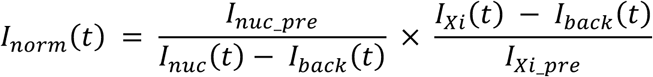

Where *I*_nuc_pre_ is the average intensity of pre-bleach nucleus intensity, *I*_Xi_pre_ is the average intensity of pre-bleach Xi intensity. Normalized intensities were then fitted to One binding state model M version^46–48^:

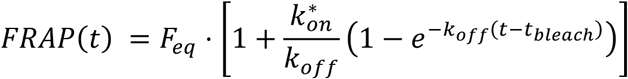

Where *F*_eq_ is the free-state fraction of transiently bound molecules, *k*_off_ is the unbinding rate, *k*^∗^ is the pseudo-binding rate constant, *t*_bleach_ is the bleach time.

For FRAP analysis of nuclear regions in male cells, images were first registered using the StackReg and TurboReg plugins. Fluorescence intensity within the bleached ROI (*I*_ROI_) was measured over time. An identically sized circular region outside the nucleus was selected to measure background fluorescence (*I*_back_) and the whole nucleus was used to measure total nuclear fluorescence (*I*^nuc^). Fluorescence intensities were normalized using the following double-normalization approach:

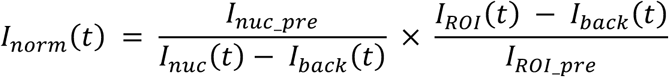

Where *I*_ROI_pre_ is the average pre-bleach intensity of bleached region. Normalized intensities were fitted to a Diffusion model for circular spot^46–48^:

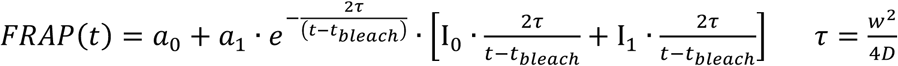

Where *a*_0_ is the unbleached fraction at bleach time *t*_bleach_, *a*_1_ is the recovery fraction, *w* is the radius of apparent bleach spot, *D* is the diffusion coefficient. Ι_0_ and Ι_1_ are modified Bessel functions. FRAP curve fitting was performed using FRAPAnalyser 2.1.0.

### Fluorescence Fluctuation Spectroscopy (FFS)

MEFs were treated with 4-OHT for 5 days and seeded into glass-bottom 8-well chamber slides (ibidi, 80827). Before imaging, cells were equilibrated in a microscope incubation chamber at 37 °C for 30 min. FFS experiments were performed on a Leica Stellaris 8 microscope as previously described, with minor modifications^53^. The Xi territory was first identified, and the brightest z-plane was selected for analysis. Live-cell line-scan imaging was performed on a Leica Stellaris 8 confocal microscope equipped with an HC PL APO CS2 63×/1.40 NA oil-immersion objective. SMCHD1-GFP was excited using a 479-nm white-light laser line. A confocal pinhole of approximately 95.6 µm, corresponding to 1 Airy unit, was used. Unidirectional x-t line scans were acquired over a ∼4.6-µm line comprising 64 spatial pixels (∼0.073 µm per pixel), with a pixel dwell time of 5.7 µs and a line acquisition time of 1 ms. Each acquisition comprised 8,192 consecutive line scans over 8.192 s and was repeated for 13 cycles, using the “maximized lines per page” setting, yielding a total acquisition time of approximately 106.5 s and 106,000 lines. Images were acquired without line or frame averaging. Scan lines were positioned to traverse the centre of the Xi territory and extend into the surrounding nucleoplasm. High-magnification reference images were acquired before and after line scanning to document the position of the scan lines relative to the region of interest. Raw intensity data were exported as TIFF files for downstream analysis.

Confocal volume calibration was performed using the Fluorescence Correlation Spectroscopy (FCS) module of the Leica Stellaris 8 microscope. Fluorescein solution (Sigma, 568864; 20 nM in 0.01 M NaOH) was equilibrated in the microscope incubation chamber at 37 °C for 30 min before measurement. The same optical settings used for line-scan FFS experiments were applied. Photon-counting measurements were acquired at a single stationary point for 1 min. Photon-count traces were transformed using the autocorrelation function (ACF):

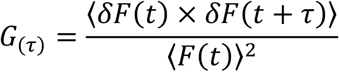

Where *F*(*t*) is the fluorescence intensity as a function of time *t*, *δF*(*t*) is the deviation of fluorescence intensity as a function of time with the respect to the mean, as *F*(*t*) − 〈*F*(*t*)〉, *δF*(*t* + *τ*) is the deviation of fluorescence intensity as a function of time with the respect to the mean shifted to every possible lag time *τ*.

Autocorrelation curve was fitted to a one-component three-dimensional diffusion model assuming a Gaussian point spread function (PSF), from which the lateral and axial radii of the confocal volume were derived:

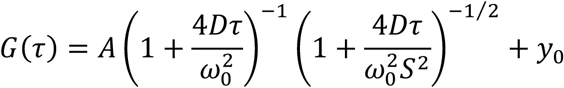

where *A* is the correlation amplitude *G*_0_, *D* is the diffusion coefficient, *τ* is the lag time, *ω*_0_ and *z*_0_ are the width and length of 3D Gaussian observation volume in the focal plane, *S* is the structure factor as *S* = *z*_0_⁄*ω*_0_, *y*_0_ is the baseline offset.

Single point FCS and line scan FFS data analysis was performed using custom MATLAB scripts based on previously described protocols, with minor modifications^53^. Raw intensity traces were corrected for photobleaching using a moving-forward detrending approach. The first 20,000 scanned lines were excluded from analysis to account for instrumental instability at the start of acquisition. The remaining pre-processed image data were used for correlation analyses. Xi, nucleoplasmic and cytoplasmic regions were identified from fluorescence-intensity carpet plots. For ACF analysis, five adjacent pixels located at the centre of the Xi territory were selected for Xi ACF fitting, and five pixels within the nucleoplasm were selected for nucleoplasmic ACF fitting. Pre-processed images were transformed using the pixel-specific ACF:

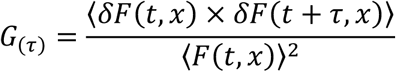

Where *F*(*t*, *x*) is the fluorescence intensity as a function of time *t* at pixel position *x*, and the rest function definitions are the same with ACF function descriptions above with pixel position limitation.

The autocorrelation curves were then fitted to a two-components 3D Gaussian diffusion model while Initial parameter estimates for ACF fitting were manually adjusted based on regional fluorescence intensities:

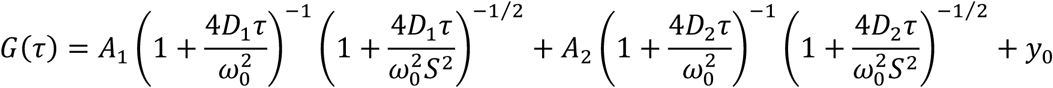

Where *D* is the diffusion coefficient, *A* is the amplitude, *ω*_0_ and *z*_0_ are the width and length of 3D Gaussian observation volume in the focal plane, *S* is the structure factor as *S* = *z*_0_⁄*ω*_0_, *y*_0_ is the baseline offset.

The multi-status fraction model was modified as described before^55^. The autocorrelation curves were fitted to model with:

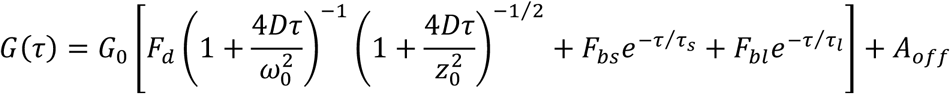

Where *G*_0_ is the autocorrelation amplitude, *D* is the diffusion coefficient, *F_d_*, *F_bs_*, and *F_bl_* are fraction of freely diffusion, short-lived bound, and long-lived bound molecules respectively. *τ_s_* and *τ_l_* are residence time of short-lived bound and long-lived bound molecules respectively. *ω*_0_ and *z*_0_ are the width and length of 3D Gaussian observation volume in the focal plane, *A_off_* is fitted baseline offset.

For pair correlation function (pCF) analysis, five pixels at the centre of the Xi territory were designated as reference Xi pixels. Pixels at defined spatial distances from the Xi reference region were selected for pCF analysis. To ensure consistency across samples, the spatial order of scan lines was manually adjusted so that directional analyses were uniformly oriented either from the Xi to nucleoplasm or from nucleoplasm to the Xi. Average signals from selected pixel positions were transformed using the pCF:

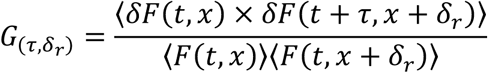

Where *F*(*t*, *x*) is fluorescence intensity as a function of time *t* and pixel position *x*, *F*(*t*, *x* + *δ_r_*) is fluorescence intensity as a function of time *t* and pixel position spatially shifted by *δ_r_* from *x*. *δF*(*t*, *x*) is the deviation of fluorescence intensity as a function of time *t* to the mean 〈*F*(*t*, *x*)〉. *δF*(*t* + *τ*, *x* + *δ_r_*) is the deviation of fluorescence intensity as a function of time *t* to the mean 〈*F*(*t*, *x* + *δ_r_*)〉 shifted by every possible lag time *τ* at pixel position *x* + *δ_r_*.

Pair correlation curves were fitted to a Gaussian model using the MATLAB fit function to derive the peak correlation and corresponding lag time. Starting values for fitting were manually adjusted to obtain biologically plausible lag times. Only samples exhibiting positive pair correlation at defined distances were retained for final pCF and mean squared displacement (MSD) analyses.

For MSD analysis, Xi or nucleoplasmic pixel regions were selected as described above. Increasing spatial offsets were then analysed (*δ_r_* = 0, 4, 8, 12, 16, 20, 24, 28 and 32 pixels) to calculate translocation lag times between positions by pCF fitting. The local diffusion coefficient *D*_0_ at each position was calculated by ACF fitting. MSD was calculated as:

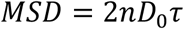

Where *n* is the dimensionality, *D*_0_ is local diffusion coefficient, *τ* is translocation lag time between selected position with each *δ_r_*.

MSD values were then fitted to an isotropic diffusion model:

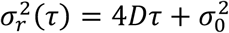

Where σ^2^*_r_*(*τ*) is the MSD, *σ*^2^ equals to *ω*^2^ which is the width of 3D Gaussian observation volume in the focal plane.

All FFS data processing and analyses were performed in MATLAB R2023b.

### Lattice light-sheet microscopy

Female NSCs were treated with 4-OHT for 5 days, and GFP-positive cells were isolated by FACS and plated in glass-bottom 8-well chamber slides (ibidi, 80827). Cells were incubated with SPY-650 DNA dye (SPIROCHROME, SC501) diluted 1:2,000 in culture medium for 1 h before imaging, according to the manufacturer’s instructions. Cells were maintained in the incubation chamber of a ZEISS Lattice Lightsheet 7 microscope at 37 °C with 5% CO_2_ and imaged using water immersion. Images were acquired as 16-bit fluorescence time series using the Sinc3SIM light-sheet mode. Light from 488 nm and 640 nm lines were directed to the sample via a 13.3 x 0.44 NA excitation objective. Resultant fluorescence was collected via a 44.83x 1.0 NA objective and projected through a multi-band quad 405/488/561/640 nm filter. For each field of view, images were acquired at 2,262 × 2,016 pixels in x-y. Z-stacks were acquired over a 50-µm axial range with 20 optical sections, and time-lapse imaging was performed for 12 h with a frame interval of approximately 132 s. Identical display and analysis settings were applied across matched experimental groups.

Maximum-intensity projections of each image series were used for downstream analysis. Xi regions were manually selected based on the number of SMCHD1-enriched foci and their nuclear localization. Xi regions were detected using the difference-of-Gaussian detector (DoG) in the TrackMate 7 FIJI plugin^89^, and temporal changes in Xi parameters, including fluorescence intensity, were recorded. During periods in which SMCHD1 Xi enrichment or nuclear localization was not visible, detection of Xi-equivalent regions was restricted to the DNA-stained area. The same Xi region size was applied for each individual cell and its daughter cells during tracking. Whole-nucleus regions were detected from the DNA staining channel using the Thresholding detector in TrackMate 7, and temporal changes in nuclear parameters, including fluorescence intensity and nuclear area, were recorded.

For each dividing cell, the time point at which SMCHD1 lost nuclear localization and dispersed into the cytoplasm was defined as time 0. All other frames were manually aligned relative to this time point. Xi GFP intensity at each time point was normalized to nuclear GFP intensity at the same time point:

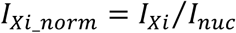

Normalized curves were then fitted to sigmoidal models^63^. The descending phase of SMCHD1 Xi enrichment was fitted to a descending Gompertz model:

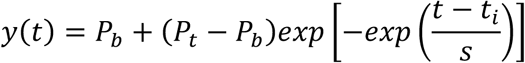

Where *P_t_* and *P_b_* are the upper and lower fitted plateaus, respectively. *t_i_* is the inflection-time parameter while *s* is descending rate constant which is the time-scale parameter controlling the steepness of decay. Practical start and endpoint times of decay were defined as the fitted times at which 95% and 5% of the fitted amplitude remained, respectively, following:

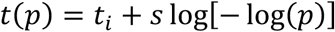

The ascending phase of SMCHD1 re-enrichment at the Xi was fitted to an ascending Gompertz model:

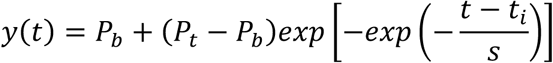

Where *P_t_* and *P_b_* are the upper and lower fitted plateaus, respectively. *t_i_* is the inflection-time parameter while *s* is descending rate constant which is the time-scale parameter controlling the steepness of decay. The practical start and endpoint of accumulation were defined as the fitted times at which 5% and 95% of the total rise were reached following:

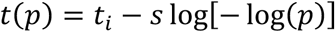

Descending fitted curves with R^2^ > 0.9 and ascending fitted curves with R^2^ > 0.8 were considered good fits and retained for analysis. All curve fitting was performed in MATLAB R2023b.

### Statistical analyses

Statistical analyses and graph generation were performed using GraphPad Prism v10.6.1 unless otherwise stated. Statistical tests were selected according to the experimental design and are specified in the corresponding figure legends. Sample sizes, numbers of independent experiments, error-bar definitions and multiple-comparison correction methods are reported in the figure legends.

## Data availability

All genomic datasets are available in the Gene Expression Omnibus under the following identifiers GSE345169 for ChIP-seq and GSE345170 for RNA-seq.

## Acknowledgements

The authors gratefully acknowledge the WEHI Centre for Dynamic Imaging, WEHI Cytometry Facility, WEHI Genomics lab, The Melbourne Advanced Genome Editing Centre (MAGEC) laboratory for their support and assistance of this work. The work was made possible via funding as follows: China Scholarship Council scholarship (RH); National Health and Medical Research Council of Australia (NHMRC) Investigator grant 1194345 (MEB), 2041117 (MEB), Investigator grant 2007996 (QG); Pamela and Lorenzo Galli Trust support (MEB); Brian M Davis Charitable Trust support (MEB); Australian Research Council (ARC) Future Fellowship FT200100401 (EH); Discovery Projects DP180101387 (EH) and DP21010298 (EH); LIEF (LE210100046, EH); and NHMRC IRIISS and Victorian State Government Operational Infrastructure Support. The generation of LSL-SMCHD1-GFP switchable mice used in this study was supported by Phenomics Australia and the Australian Government through the National Collaborative Research Infrastructure Strategy (NCRIS) program.

## Author Contributions

Conceptualization: MEB, EH, RH; Methodology: MEB, RH, EH, NDG, JSV, ANS, JL, AKe, TW; Validation: RH, TC, KB; Formal Analysis: RH, JSV, JL, Ake; Investigation: RH, JL, KB, TC, IW, QG, TW; Resources: MEB, AKu; Writing – Original Draft: RH; Writing – Review and Editing: RH, MEB, EH, NDG; Visualisation: RH; Supervision: MEB, EH, NDG; Project Administration: MEB; Funding Acquisition: MEB, RH, QG, EH

