## Supplementary figures for "SMCHD1’s DNA binding activity enables its stable retention on chromatin"

### Extended Data

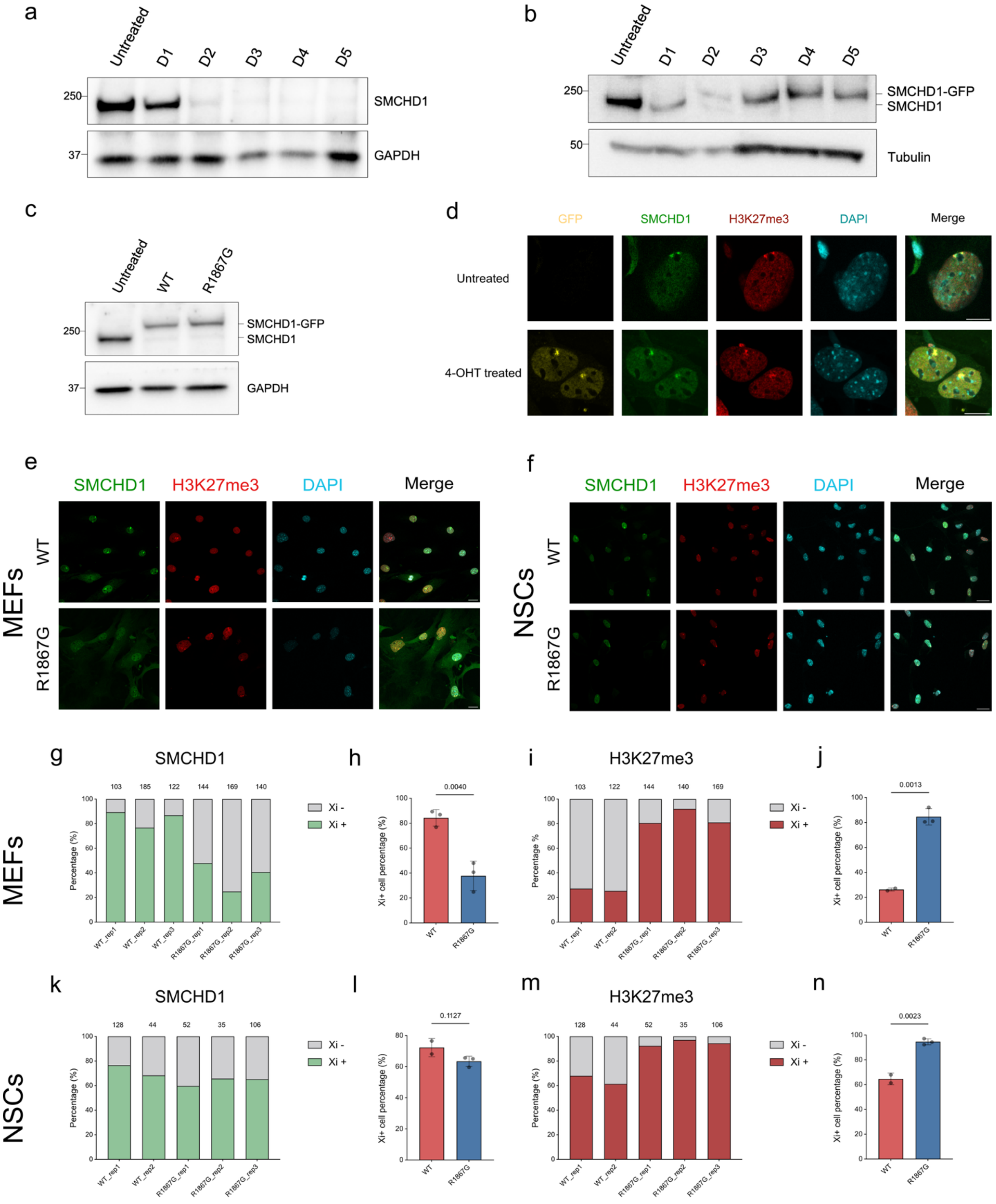

**Extended Data Fig. 1 Validation of switchable SMCHD1 cell lines and impaired SMCHD1 Xi-domain formation following disruption of DNA binding.** a, Immunoblot analysis of endogenous SMCHD1 depletion after 1-5 days of 4-OHT treatment in an Smchd1 conditional-knockout NSC line (*Smchd1*$^{fl/fl}$; *Rosa26*$^{CreERT2/+)}$. b, Immunoblot analysis of simultaneous endogenous SMCHD1 depletion and SMCHD1-GFP induction after 1-5 days of 4-OHT treatment in a switchable NSC line (*Smchd1*$^{fl/fl}$; *Rosa26*$^{LSL\text{-}SMCHD1\text{-}GFP/CreERT2}$). c, Immunoblot analysis of WT and R1867G SMCHD1-GFP expression in NSCs after 5 days of 4-OHT treatment. Blots were probed with antibodies against SMCHD1, tubulin or GAPDH. d, Representative immunofluorescence images showing SMCHD1-GFP enrichment at the Xi in MEFs after 5 days of 4-OHT treatment. Cells were stained with antibodies against GFP, SMCHD1 and H3K27me3 and counterstained with DAPI. Scale bar, 10 μm. e, Representative immunofluorescence images of 4-OHT-treated MEFs. Scale bar, 20 μm. f-i, Percentage of MEFs with a detectable SMCHD1- or H3K27me3-enriched Xi domain. j, Representative immunofluorescence images of 4-OHT-treated NSCs. Scale bar, 20 μm. k-n, Percentage of NSCs with a detectable SMCHD1- or H3K27me3-enriched Xi domain. Data in f-i and k-n were pooled from independent experiments, and P values were calculated using two-sided t-tests. The number of cells analysed in each independent experiment is indicated above the corresponding data point.

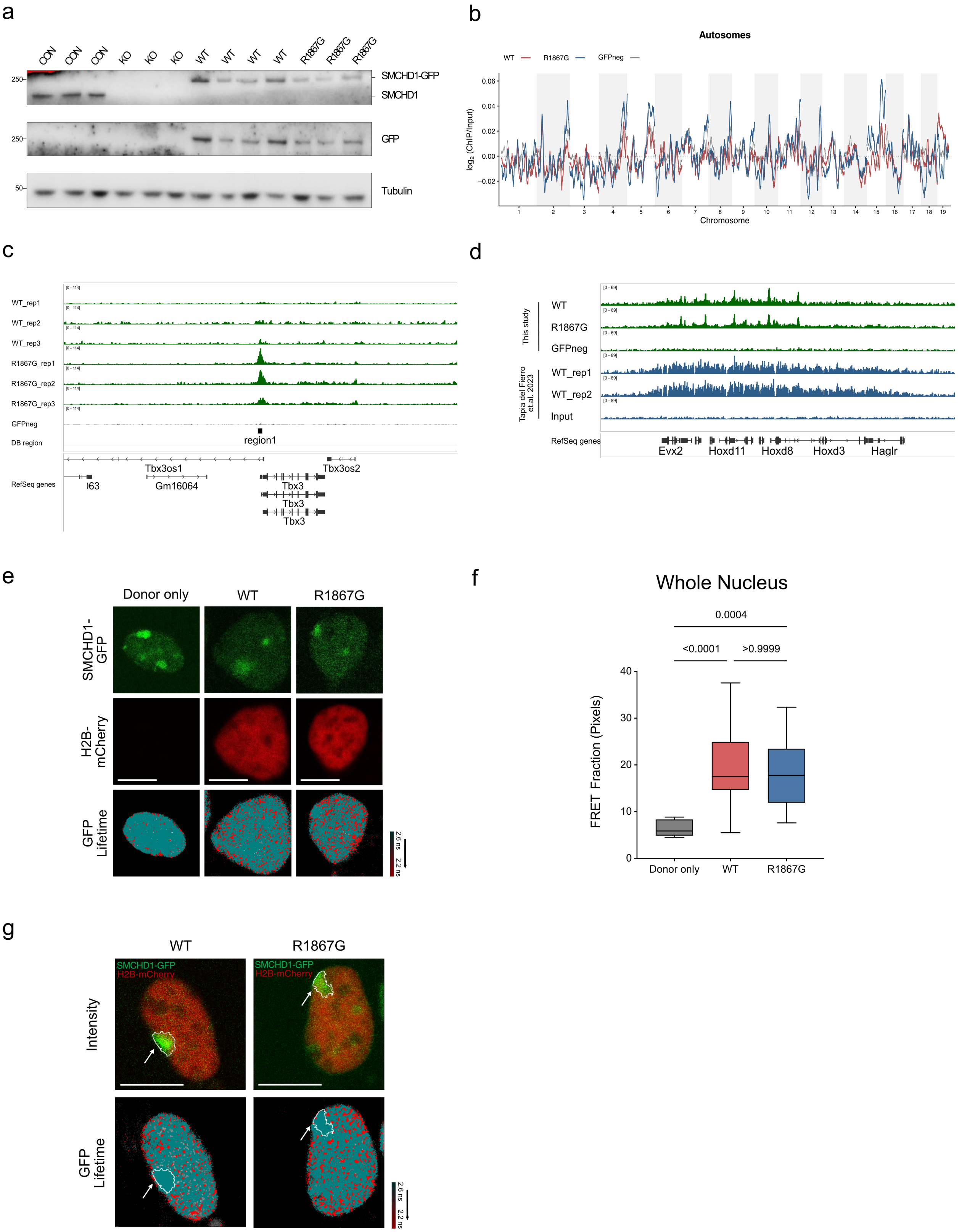
a
CON
CON
CON
KO
KO
KO
WT
WT
WT
WT
R1867G
R1867G
R1867G
250
SMCHD1-GFP
SMCHD1
250
GFP
50
Tubulin
b
Autosomes
WT
R1867G
GFPneg
0.06
0.04
0.02
0.00
-0.02
log2 (ChIP/Input)
Chromosome
c
WT_rep1
WT_rep2
WT_rep3
R1867G_rep1
R1867G_rep2
R1867G_rep3
GFPneg
DB region
region1
RefSeq genes
Tbx3os1
Gm16064
Tbx3
Tbx3
Tbx3
Tbx3os2
d
This study
WT
R1867G
GFPneg
Tapia del Fierro et.al. 2023
WT_rep1
WT_rep2
Input
RefSeq genes
Evx2
Hoxd11
Hoxd8
Hoxd3
Haglr
e
Donor only
WT
R1867G
SMCHD1-GFP
H2B-mCherry
GFP Lifetime
2.6 ns
2.2 ns
f
Whole Nucleus
0.0004
<0.0001
>0.9999
FRET Fraction (Pixels)
40
30
20
10
0
Donor only
WT
R1867G
g
WT
R1867G
SMCHD1-GFP
H2B-mCherry
Intensity
GFP Lifetime
2.6 ns
2.2 ns

Extended Data Fig. 2 Validation of SMCHD1 expression and autosomal ChIP-seq profiles in switchable NSCs. a, Immunoblot analysis of endogenous SMCHD1 and induced SMCHD1-GFP in female NSCs after 4-OHT treatment. Genotypes were as follows: control (CON), *Smchd1*$^{fl/fl}$; *Rosa26*$^{+/+}$; knockout (KO), *Smchd1*$^{fl/fl}$; *Rosa26*$^{CreERT2/+;}$ WT and R1867G, *Smchd1*$^{fl/fl}$; *Rosa26*$^{LSL\text{-}SMCHD1\text{-}GFP/CreERT2}$. Blots were probed with antibodies against SMCHD1, GFP and tubulin. b, SMCHD1 ChIP-seq profiles across autosomes. For clarity, only group means are shown. Bin size, 10 Mb. c, Differential binding of SMCHD1 R1867G mutant at *Tbx3* locus. d, SMCHD1 occupancy at the *HoxD* locus detected using the indicated ChIP-seq strategies. e, Representative images show interaction between SMCHD1 and histone H2B in live female NSCs. Top panel: NSCs expressing SMCHD1-GFP; Middle panel: NSCs transfected with or without H2B-mCherry; Bottom panel: pseudocolored GFP lifetime maps of cells. Scale bar, 5 μm. f, Fraction of pixels that detected whole-nucleus FRET signal of each group in live female NSCs. Combined samples of two independent experiments. n = 8 Donor only, 21 WT and 23 R1867G cells. P values were calculated by one-way ANOVA with Bonferroni correction for multiple comparisons. g, Representative images show compromised FLIM-FRET detection at the Xi territory in live female NSCs. The Xi territory was highlighted based on SMCHD1-GFP intensity threshold. Top panel: intensity images of cell expressing SMCGD1-GFP (green) and H2B-mCherry (red); Bottom panel: pseudocolored GFP lifetime maps of cells. Scale bar, 5 μm.

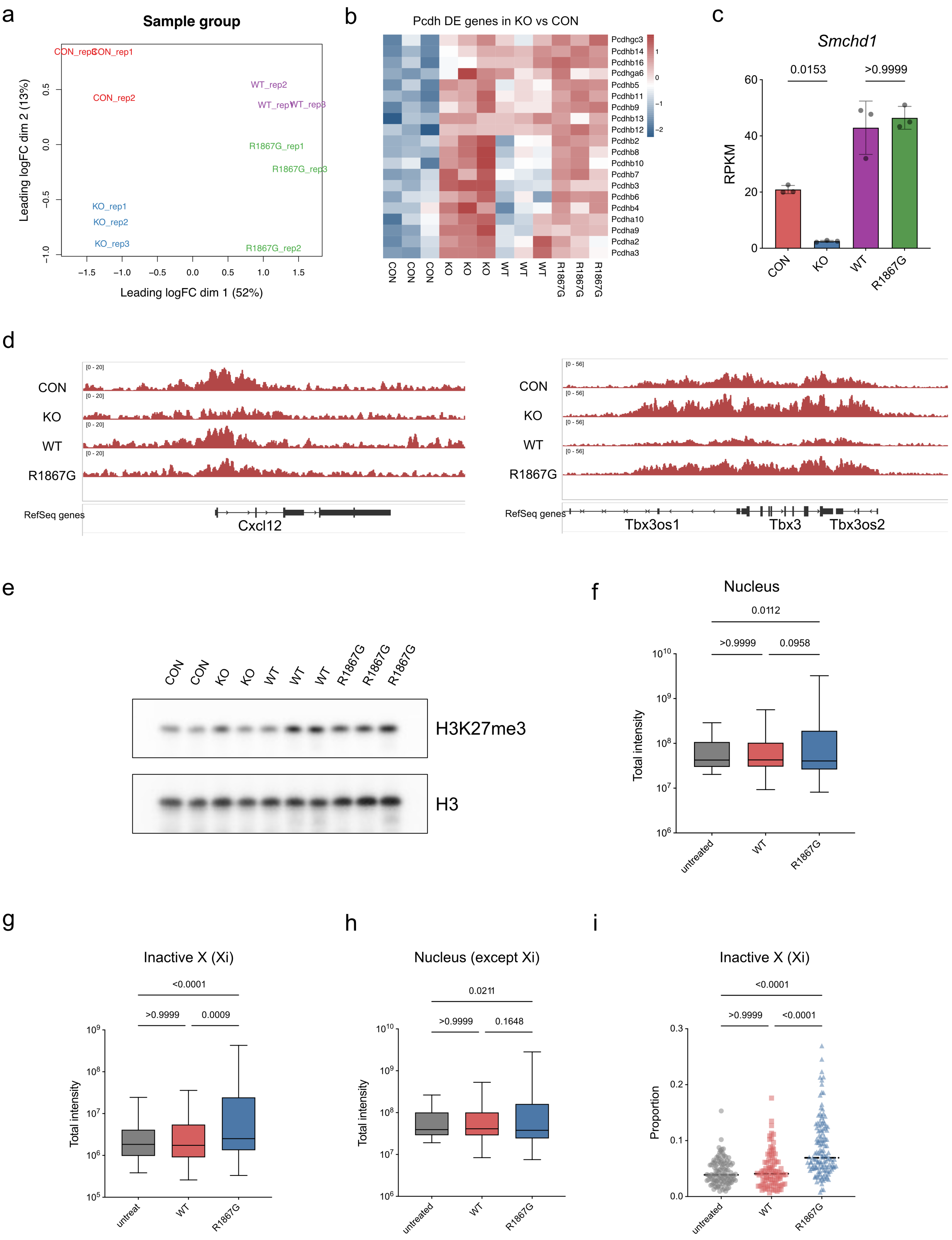
a
Sample group
Leading logFC dim 1 (52%)
Leading logFC dim 2 (13%)
CON_rep1
CON_rep2
CON_rep3
WT_rep1
WT_rep2
WT_rep3
R1867G_rep1
R1867G_rep2
R1867G_rep3
KO_rep1
KO_rep2
KO_rep3
b
Pcdh DE genes in KO vs CON
Pcdhgc3
Pcdhb14
Pcdhb16
Pcdhga6
Pcdhb5
Pcdhb11
Pcdhb9
Pcdhb13
Pcdhb12
Pcdhb2
Pcdhb8
Pcdhb10
Pcdhb7
Pcdhb3
Pcdhb6
Pcdhb4
Pcdha10
Pcdha9
Pcdha2
Pcdha3
c
Smchd1
RPKM
0.0153
>0.9999
CON
KO
WT
R1867G
d
CON
KO
WT
R1867G
RefSeq genes
Cxcl12
Tbx3os1
Tbx3
Tbx3os2
e
H3K27me3
H3
f
Nucleus
Total intensity
0.0112
>0.9999
0.0958
untreated
g
Inactive X (Xi)
<0.0001
0.0009
untreat
h
Nucleus (except Xi)
0.0211
0.1648
i
Inactive X (Xi)
Proportion
<0.0001

**Extended Data Fig. 3 R1867G is a hypomorphic SMCHD1 variant that redistributes H3K27me3 without altering global abundance.** a, Multidimensional-scaling plot of RNA-seq samples. b, Expression of *Pcdh* genes identified as differentially expressed in SMCHD1-KO cells across CON, KO, WT and R1867G NSCs. c, Smchd1 expression across RNA-seq groups. P values were calculated by one-way ANOVA. d, H3K27me3 occupancy at the *Cxcl12* and *Tbx3* loci, where R1867G and KO cells show concordant changes relative to their respective controls. e, Immunoblot analysis of total H3K27me3 in the NSC lines used for ChIP-seq. f, Total H3K27me3 fluorescence intensity within the nuclei of NSCs analysed in Fig. 1. g,h, Total H3K27me3 fluorescence intensity within the Xi and within the remainder of the nucleus, respectively. The same three-dimensional Xi masks used in Fig. 1 were applied. i, Fraction of H3K27me3 fluorescence within the Xi to the whole nucleus ($I_{\text{xi}}$ / $I_{\text{Total Nucleus}}$). For f-i, P values were calculated by one-way ANOVA with Bonferroni correction for multiple comparisons.

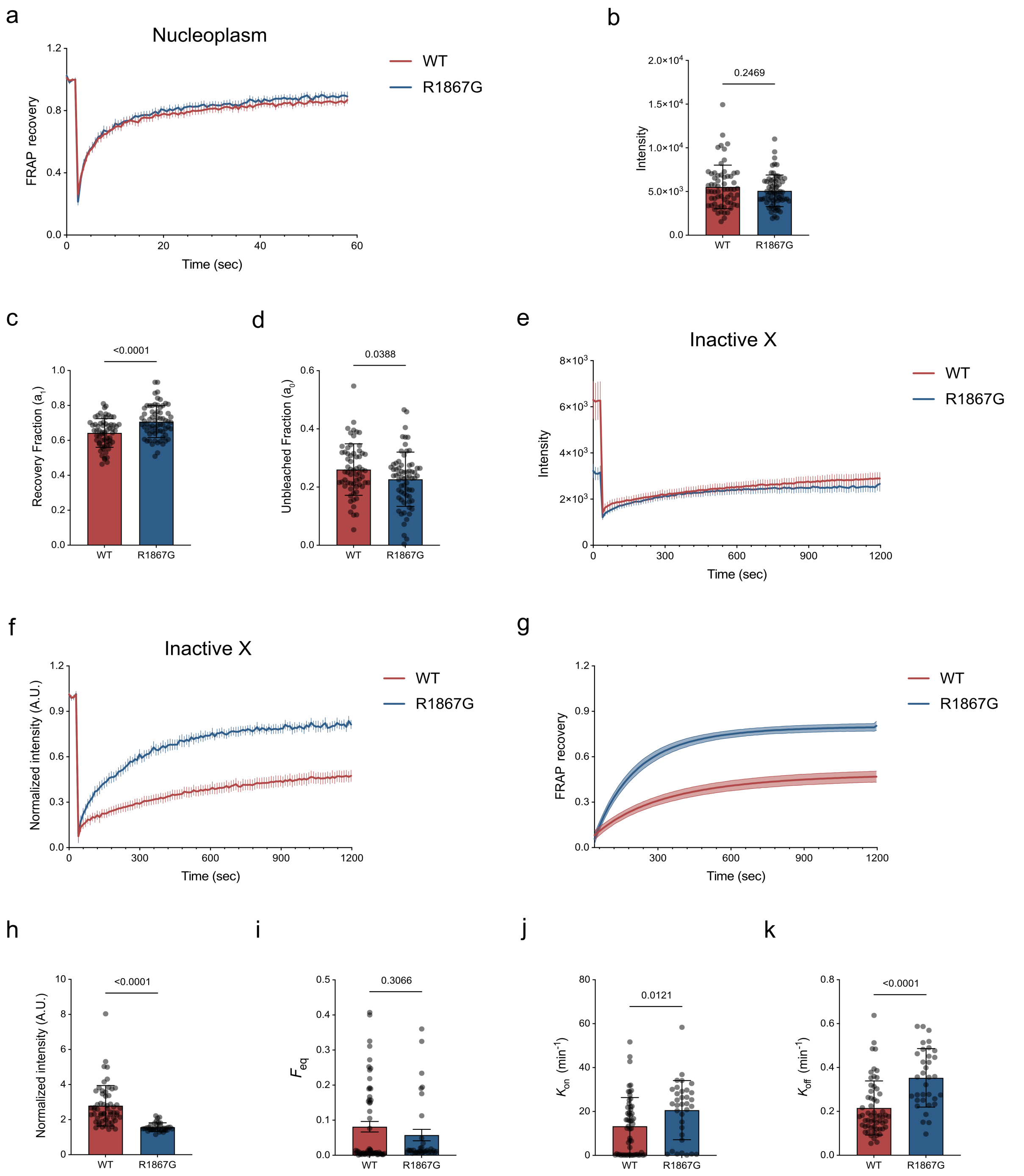


**Extended Data Fig. 4 FRAP dynamics of WT and R1867G SMCHD1 in unselected cell populations.** a, Normalized nucleoplasmic FRAP recovery curves in male MEFs. Data are mean with 95% confidence intervals. b, Mean pre-bleach fluorescence across the entire nucleus of male MEFs. Data are mean ± s.d.; the P value was calculated using a two-sided t-test. c, Recovery fraction ($\alpha_1$) of SMCHD1 fluorescence in observed time window after bleaching. d, Unbleached fraction ($\alpha_0$) of SMCHD1 fluorescence at the bleaching timepoint. For c and d, combined results from 3 independent FRAP experiments in male MEFs. Data are mean ± s.d.; the P value was calculated using a two-sided t-test. n = 63 WT and 67 R1867G cells. e, Unnormalized Xi fluorescence before and after bleaching in all female MEFs. f, Normalized Xi FRAP recovery curves in all female MEFs. g, Fitted Xi FRAP recovery curves in all female MEFs. In e-g, data are mean with 95% confidence intervals. h, Normalized pre-bleach Xi fluorescence in all cells. i, Freely diffusing fraction of SMCHD1 at the Xi. j, Apparent association rate of freely diffusing SMCHD1 with chromatin. k, Dissociation rate of chromatin-bound SMCHD1. For h-k, data were pooled from three independent experiments; n = 58 WT and 34 R1867G cells. Data in h,j,k are mean ± s.d.; data in i is mean ± s.e.m. P values were calculated using two-sided t-tests.

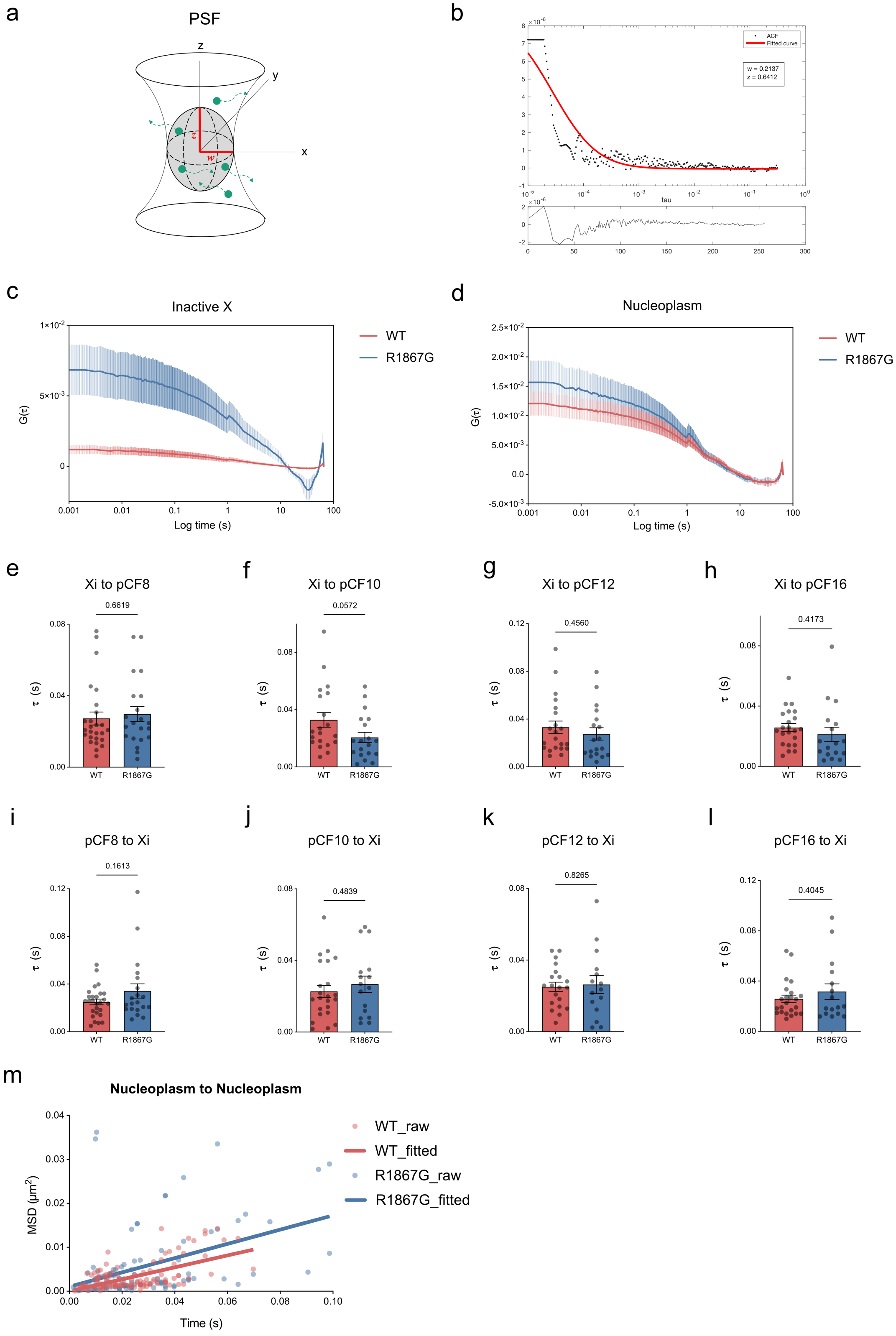

a
PSF
b
ACF
Fitted curve
w = 0.2137
z = 0.6412
tau
c
Inactive X
WT
R1867G
G(τ)
Log time (s)
d
Nucleoplasm
WT
R1867G
G(τ)
Log time (s)
e
Xi to pCF8
0.6619
f
Xi to pCF10
0.0572
g
Xi to pCF12
0.4560
h
Xi to pCF16
0.4173
i
pCF8 to Xi
0.1613
j
pCF10 to Xi
0.4839
k
pCF12 to Xi
0.8265
l
pCF16 to Xi
0.4045
τ (s)
WT
R1867G
m
Nucleoplasm to Nucleoplasm
WT_raw
WT_fitted
R1867G_raw
R1867G_fitted
MSD (μm²)
Time (s)

**Extended Data Fig. 5 Confocal calibration and distance-resolved pCF analysis of SMCHD1 exchange between the Xi and the nucleoplasm.** a, Schematic of the three-dimensional confocal detection volume (grey oval). The lateral radius ($\omega$) and axial radius ($z$) define a Gaussian detection volume governed by the point spread function (PSF). Green dots denote fluorescent molecules. b, Calibration of the confocal detection volume using fluorescein. The autocorrelation curve was fitted using a one-component three-dimensional diffusion model. c,d, Raw ACF curves within the Xi (c) and the nucleoplasm (d). e-h, Arrival times calculated from positions within the Xi to nucleoplasmic positions at the indicated offsets. The pixel size was 73.22 nm. i-l, Arrival times calculated from nucleoplasmic positions to positions within the Xi at the indicated offsets. For e-l, data are mean ± s.e.m. and P values were calculated using two-sided t-tests. m, Mean-squared-displacement analysis of long-range SMCHD1 movement between two nucleoplasmic positions. Directional diffusion coefficients: WT, 0.034 $\mu m^2 \cdot s^{-1}$; R1867G, 0.041 $\mu m^2 \cdot s^{-1}$.

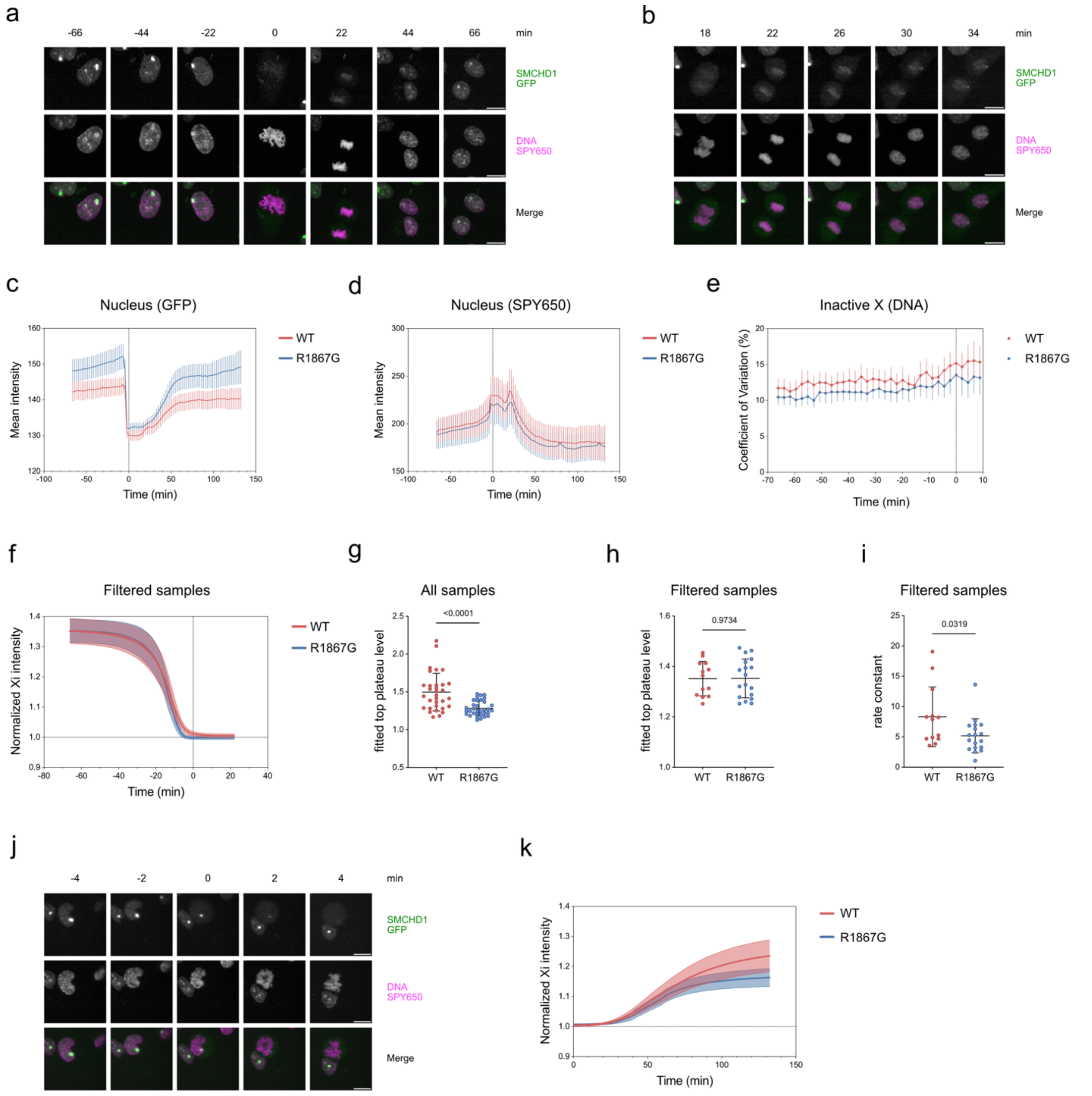


Extended Data Fig. 6 SMCHD1 is released from the Xi at defined stages of mitosis. a, Representative live-cell images showing loss of Xi-associated SMCHD1 during mitosis and re-accumulation in daughter cells. Scale bar, 10 µm. b, Representative live-cell images showing recruitment of SMCHD1 to the Xi during cytokinesis. Scale bar, 10 µm. c, Mean SMCHD1-GFP fluorescence within the nuclear region during mitosis. The nuclear region was defined using the DNA-staining channel. Data are mean with 95% confidence intervals. d, Mean DNA-staining intensity within the nuclear region during mitosis. Data are mean with 95% confidence intervals. e, Coefficient of variation of DNA fluorescence within the Xi, calculated as s.d./mean × 100%. Xi was defined using the strongest SMCHD1-GFP Xi domain, and the same mask size was applied across time points. Data are mean with 95% confidence intervals. f, Fitted dissociation curves for WT and R1867G cells selected for comparable pre-mitotic Xi fluorescence. Data are mean with 95% confidence intervals; n = 13 WT and 18 R1867G cells. g, Fitted upper plateau of all dissociation curves shown in Fig. 6c. h, Fitted upper plateau for the intensity-matched cells shown in f. i, Dissociation-rate constant for the intensity-matched cells. In g-i, data are mean ± s.d. and P values were calculated using two-sided t-tests. j, Representative live-cell images showing retention of SMCHD1 at the Xi after nuclear envelope breakdown timepoint in a subset of cells. Scale bar, 10 μm. k, Fitted accumulation curves for all WT and R1867G cells. Data are mean with 95% confidence intervals; n = 36 WT and 33 R1867G cells. Data were pooled from two independent experiments.

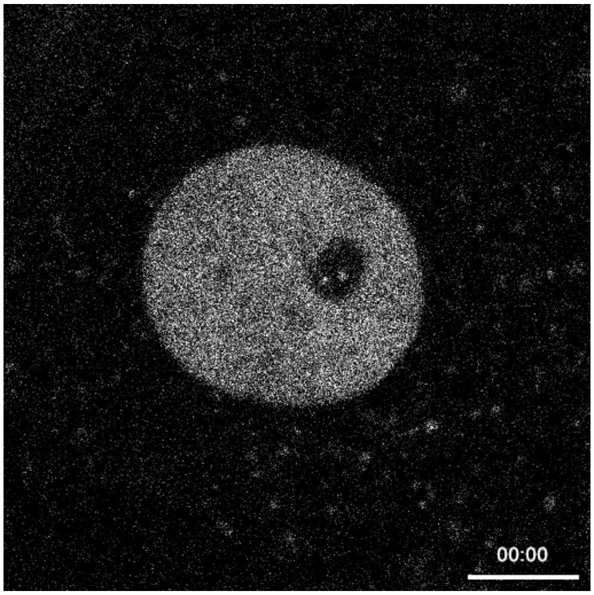


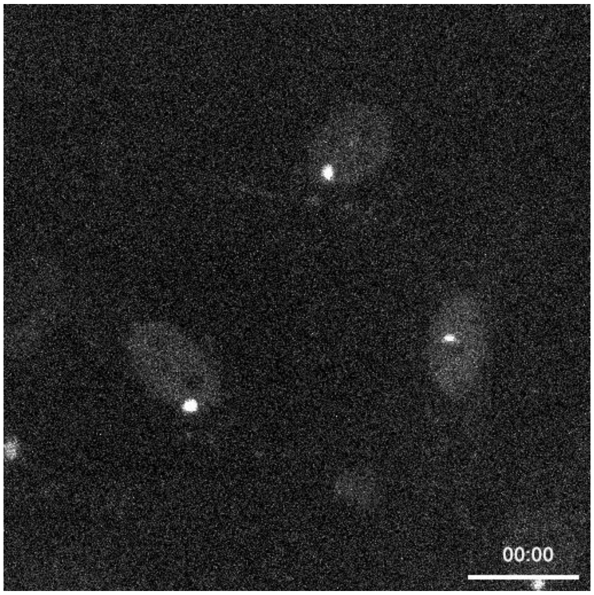


Supplementary Video 1. Left, representative nucleoplasmic FRAP experiment in a male MEF. Scale bar, 10 μm.

Supplementary Video 2. Right, representative Xi FRAP experiment in a female MEF. Scale bar, 20 μm.

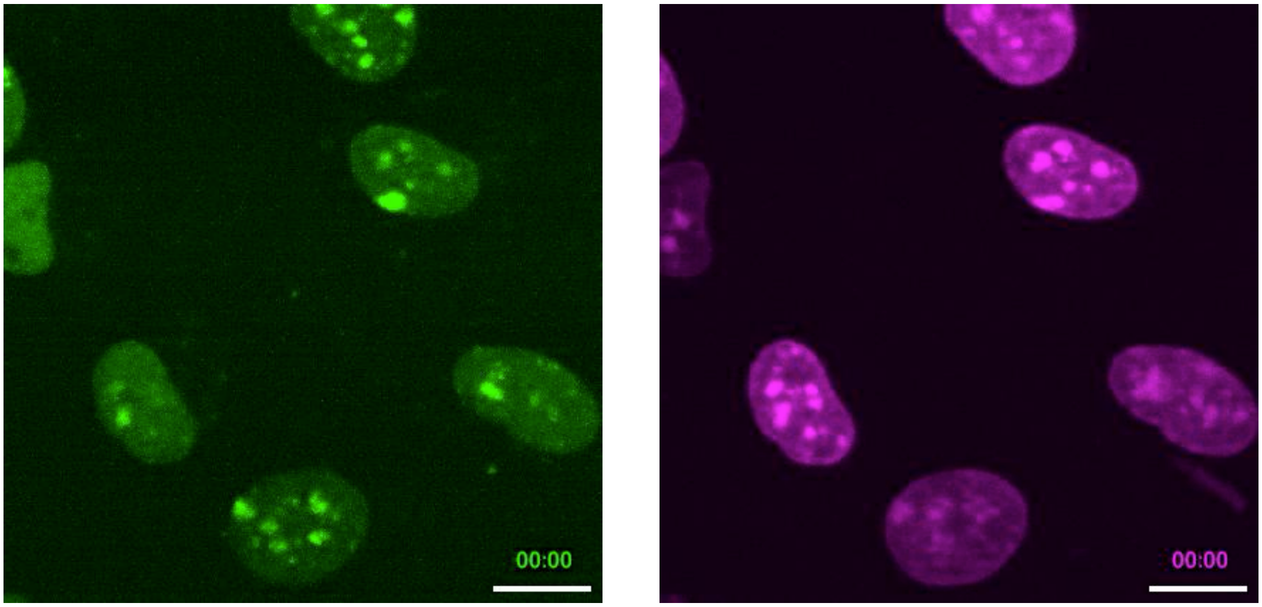


Supplementary Video 3: Example of SMCHD1 dynamics during mitosis in female NSCs. Left: GFP channel of NSCs expressing SMCHD1-GFP. Right: SPY-650 DNA dye stained NSCs. Scale bar represents 10μm.
