## Supplementary tables 1 and 2 for "SMCHD1’s DNA binding activity enables its stable retention on chromatin"

Supplementary Table 1. Genotyping primers

| **Target** | **Name** | **Sequence** |
| --- | --- | --- |
| X chromosome | Otc F | GTTCTTTCGTTTTCCCCTCTC |
| X chromosome | Otc R | GGCATTATCTAAGGAGGAGCATC |
| Y chromosome | Zfy F | GACTAGACATGTCTTAACATCTGTCC |
| Y chromosome | Zfy R | CCTATTGCATGGACAGCAGCTTATG |
| Endogenous SMCHD1 KO | SMCHD1 CKO F2 | TCAGGTGGTCTCGAGCCC |
| Endogenous SMCHD1 KO | SMCHD1 CKO F4 | CCATGAGAAGCAATGTGGGA |
| Endogenous SMCHD1 KO | SMCHD1 CKO R1 | GGACAGCCAAAGTGACACAG |
| CreERT2 knock-in | WA 693 | CTGACCGTACACCAAAATTTGCCTG |
| CreERT2 knock-in | WA 694 | GATAATCGCGAACATCTTCAGGTTC |
| LSL-SMCHD1 knock-in | CTV SMCHD1 F1 | GCCTCCTGGCTTCTGAGGACCG |
| LSL-SMCHD1 knock-in | CTV SMCHD1 F2 | TCTGTGGGAAGTCTTGTCCCTCC |
| LSL-SMCHD1 knock-in | CTV SMCHD1 R1 | CCTGGACTACTGCGCCCTACAGA |
| LSL-SMCHD1 KI (long) | CTV SMCHD1 long F1 | ACACTGGAATGGATCTGGTTGGCACTGTAG |
| LSL-SMCHD1 KI (long) | CTV SMCHD1 long R1 | TGTTGATCTCGAAGACCTGTTGCTGCTCAG |

Supplementary Table 2. Antibodies

| **Antibody** | **Manufacturer** | **Cat. Number** | **Host Species** | **Usage** |
| --- | --- | --- | --- | --- |
| SMCHD1 | WEHI in-house  antibody #8 /  Merck-Millipore | MABS2292 | Rat | 1:100 (IF)  1:500 (WB) |
| GFP | Invitrogen | A11122 | Rabbit | 1:1000 (WB)  10 μg (ChIP) |
| GFP | Abcam | ab13970 | Chicken | 1:100 (IF) |
| H3K27me3-  Alexa 647 | Cell Signalling  Technologies | C36B11 (#12158) | Rabbit | 1:100 (IF) |
| H3K27me3 | Cell Signalling  Technologies | 9733S | Rabbit | 1:1000 (WB)  2 μg (ChIP) |
| Spike-in | Active Motif | 104597 | Mouse | 1 μg (ChIP) |
| GAPDH | Cell Signalling  Technologies | 14C10 | Rabbit | 1:5000 (WB) |
| Tubulin | Merck | T9026 | Mouse | 1:5000 (WB) |
| H3 | Abcam | ab1791 | Rabbit | 1:2000 (WB) |
| Alexa Flour  Anti-Mouse 488 | Invitrogen | A21102 | Donkey | 1:500 (IF) |
| Alexa Flour  Anti-Mouse 546 | Invitrogen | A11040 | Goat | 1:500 (IF) |
